# Stable eusociality via maternal manipulation when resistance is costless

**DOI:** 10.1101/019877

**Authors:** Mauricio González-Forero

## Abstract

In many eusocial species, queens use pheromones to influence offspring to express worker phenotypes. While evidence suggests that queen pheromones are honest signals of the queen’s reproductive health, here I show that queen’s honest signaling can result from ancestral maternal manipulation. I develop a mathematical model to study the coevolution of maternal manipulation, offspring resistance to manipulation, and maternal resource allocation. I assume that (1) maternal manipulation causes offspring to be workers against offspring’s interests; (2) offspring can resist at no direct cost, as is thought to be the case with pheromonal manipulation; and (3) the mother chooses how much resource to allocate to fertility and maternal care. In the coevolution of these traits, I find that maternal care decreases, thereby increasing the benefit that offspring obtain from help, which in the long run eliminates selection for resistance. Consequently, ancestral maternal manipulation yields stable eusociality despite costless resistance. Additionally, ancestral manipulation in the long run becomes honest signaling that induces offspring to help. These results indicate that both eusociality and its commonly associated queen honest signaling can be likely to originate from ancestral manipulation.

## Introduction

Eusocial organisms form colonies that are distinctly influenced by their queens. In many species, a eusocial colony is composed of one queen and largely non-reproductive workers that are the queen’s offspring (Wilson, 1971, Michener, 1974). Whether a queen’s offspring becomes a worker or a future queen is often mediated by the queen herself: for example, (1) the queen in some social wasps and bees maintains the reproductive monopoly of the colony through aggression (Fletcher and Ross, 1985); (2) in many social insects the queen can feed offspring with food of different quantity or quality influencing offspring’s future reproductive caste (i.e., queen or worker) (e.g., O’Donnell, 1998, Bourke and Ratnieks, 1999, Kapheim *et al.*, 2011, Brand and Chapuisat, 2012); (3) in an ant species the queen can deposit hormones in the eggs inducing offspring to develop into workers (Schwander *et al.*, 2008); (4) in certain wasp and termite species the queen can produce pheromones that prevent offspring from becoming queens (Bhadra *et al.*, 2010, Matsuura *et al.*, 2010); and (5) in honeybees queen pheromones can induce workers to feed larvae without royal jelly causing larvae to develop into workers (Le Conte and Hefetz, 2008, Kamakura, 2011). In addition to influencing caste determination, queens can use pheromones to keep workers’ ovaries undeveloped (e.g., Holman *et al.*, 2010, Van Oystaeyen *et al.*, 2014), and to alter workers’ brain functioning inducing workers to perform various tasks (Beggs *et al.*, 2007). Although other factors can influence offspring’s worker phenotype (e.g., environmental temperature, colony size, colony age, and offspring genetic predisposition; Lo *et al.*, 2009, Schwander *et al.*, 2010), queen influence on worker development, sterility, and behavior is widespread in eusocial taxa (Fletcher and Ross, 1985, Le Conte and Hefetz, 2008, Schwander *et al.*, 2010).

The function of queen influence is typically interpreted in terms of either manipulation or honest signaling (Dawkins and Krebs, 1978, Keller and Nonacs, 1993). Manipulation refers to altering a recipient individual’s phenotype against its inclusive fitness interests (Dawkins, 1978, 1982), as is increasingly well documented in host manipulation by parasites (Poulin, 2010, Maure *et al.*, 2011, 2013, Dheilly *et al.*, 2015). In contrast, signaling refers to altering a recipient’s phenotype in its inclusive fitness interests, provided that the signaler evolved to produce that effect and the recipient to attend the signal (Maynard Smith and Harper, 2003). Manipulation and honest signaling thus differ in that the former implies conflict while the latter does not.

The presence or absence of conflict entails contrasting evolutionary patterns. On the one hand, manipulation by the queen implies that the population can be in one of three possible stages: in an ongoing arms race between manipulation and resistance to it, in successful manipulation if resistance is costly enough, or in successful resistance if resistance is sufficiently cost-free (e.g., Trivers, 1974, Craig, 1979, Uller and Pen, 2011, González-Forero and Gavrilets, 2013). On the other hand, queen honest signaling implies mutually beneficial coevolution of queen influence and offspring response (Keller and Nonacs, 1993, Maynard Smith and Harper, 2003). Then, a key factor allowing to distinguish manipulation from honest signaling is the cost of resistance: if resistance is rather costless and no arms race is detected, queen influence is expected to more likely be honest signaling (Keller and Nonacs, 1993). In particular, queen influence via pheromones is thought to be rather costless to resist and is thus considered more likely to be honest signaling than manipulation (Keller and Nonacs, 1993) as is increasingly supported by the evidence (e.g., Heinze and d’Ettorre, 2009, van Zweden *et al.*, 2014).

Then, while the commonality of queen influence has long suggested that eusociality can be caused by maternal manipulation (Alexander, 1974, Michener and Brothers, 1974, Linksvayer and Wade, 2005, Russell and Lummaa, 2009), this has not been supported by the evidence of queen honest signaling. Here I describe a mechanism that offers an explanation for the lack of evidence of manipulation. In this mechanism, maternal manipulation yields eusociality while becoming honest signaling in the long run.

Manipulation is of particular interest because it allows eusociality to evolve under relatively lax conditions when resistance cannot evolve (e.g., Trivers, 1974, Charlesworth, 1978). Without maternal manipulation, the genes for helping behavior are in the offspring who then control their own behavior. Under standard assumptions, helping is then favored when the fitness cost to the helper (*c*) is smaller than the fitness benefit to the recipient (*b*) weighted by their relatedness (*r*; i.e., *br* > *c*, Hamilton, 1964, Frank, 1998). In contrast, with maternal manipulation and disregarding resistance, the genes for helping are by definition in the mother, who then controls offspring helping behavior. Helping is in this case favored under smaller benefit-cost ratios (e.g., *b*/*c* > 1 rather than *b*/*c* > 1/*r*), because the costs of helping are paid by the helper rather than by the individual controlling the behavior (e.g., Trivers, 1974, Charlesworth, 1978). Now, if resistance can occur but is costly enough to be disfavored, manipulation is still particularly likely to generate eusociality because of the smaller benefit-cost ratios required (González-Forero and Gavrilets, 2013).

Yet, if manipulation can be resisted at no cost, the evolution of offspring resistance is expected to destabilize eusociality (Trivers, 1974, Craig, 1979, Keller and Nonacs, 1993, Uller and Pen, 2011). This view is suggested by a variety of relevant mathematical models of evolutionary conflict (Ratnieks, 1988, Ratnieks and Reeve, 1992, Reeve and Keller, 2001, Wenseleers *et al.*, 2003, 2004b,a, Cant, 2006, Ratnieks *et al.*, 2006, Shen and Reeve, 2010, Uller and Pen, 2011, Dobata, 2012, González-Forero and Gavrilets, 2013, González-Forero, 2014). A possible way to stabilize eusociality via manipulation is suggested by a study where the evolution of the benefit eliminates the mother-offspring conflict over helping behavior (González-Forero, 2014). Still, such disappearance of conflict requires that a form of resistance is costly (i.e., helping inefficiency). Since resistance to queen pheromones is presumably costless, it is of particular interest to determine if eusociality can be stabilized even when there are no direct costs associated with resistance.

With this aim, I develop a model for the coevolution of maternal manipulation, offspring costless resistance, and maternal resource allocation into fertility and maternal care. I show that the coevolution of these traits yields a reduction of maternal care that increases the benefit that offspring receive from help. This eliminates the mother-offspring conflict over helping behavior and stabilizes eusociality. These results rely on the assumption that offspring receiving no maternal care use help more efficiently than offspring receiving maternal care. In contrast to previous findings, this form of conflict resolution can occur without any direct costs of resistance.

## Model

### Key assumptions

I consider a population with parental care. For concreteness, I take parental care to be brood provisioning, although it can be any form of parental care directed to individual offspring rather than to an entire brood (e.g., some forms of brood defense; Cocroft, 2002). Each mother produces and provisions two subsequent broods, and then dies. The first brood reaches adulthood while the second one is not yet mature, so generations are overlapping. This form of reproduction is common in primitively eusocial paper wasps and sweat bees as well as in their solitary sister taxa (Michener, 1990, Hunt, 2007). Upon reaching adulthood, all adults disperse from their natal nest to a common mating pool. All individuals in the mating pool mate once and randomly. This assumption of single mating follows the evidence that monogamy is ancestral to eusociality (Hughes *et al.*, 2008, Boomsma, 2009). After mating, females compete globally for patches with resources to establish their nests. Each successful female secures a patch with resources and allocates the secured resources into producing and provisioning offspring of the two broods, closing the life cycle.

I study the coevolution of five traits: one for maternal influence, one for offspring resistance, and three describing maternal resource allocation to fertility and care of the two broods. Maternal influence is a trait that allows the mother to influence first-brood offspring to stay in the natal nest as adults (e.g., by disrupting the physiological process that urges offspring to leave, say by means of a pheromone). Maternal influence is thus a maternal effect trait (Wolf and Wade, 2009). Influenced offspring can acquiesce (i.e., not resist) by staying as adults in their natal nest and by expressing some of their usual parental care behaviors. A similar form of acquiescence is known in hosts that are manipulated by parasites to perform defense behaviors (Maure *et al.*, 2011, 2013). The parental care behaviors expressed by acquiescing first-brood offspring are received by the available brood which are second-brood offspring (i.e., helping is directed toward full siblings). I refer to an acquiescing individual as a helper. If a second-brood offspring receives help, its survival increases, where offspring survival is defined as the probability to become a parent.

Alternatively, the offspring resistance trait allows influenced offspring to resist the maternal influence by leaving the nest to mate without delay and without incurring any direct fitness loss (e.g., by reducing the number of binding site receptors of the pheromone; as discussed by Kuijper and Hoyle, 2015). Similar dispersal behaviors are known for first-brood individuals leaving their natal nest in primitively eusocial paper wasps (Reeve *et al.*, 1998) and sweat bees (Yanega, 1988). I assume the effectiveness of resistance to maternal influence to be weak at the start of the coevolutionary process because individuals have not been previously exposed to such maternal influence. An analogous example of weak resistance has been experimentally documented in microorganisms when exposed to novel parasites (Lohse *et al.*, 2006).

Maternal resource allocation occurs as follows. One trait describes how much resource the mother devotes to each of the two broods, and the other two traits (one for each brood) describe how much of this resource she spends in producing and provisioning offspring. The three maternal resource allocation traits are controlled by the mother. An offspring is either provisioned or not by the mother, and I refer to the former as maternally provisioned and to the latter as maternally neglected. These two properties describe an offspring condition. After the mother has had the opportunity to provision offspring, both maternally provisioned and neglected offspring can be provisioned by helpers. I refer to offspring provisioned by helpers as helped offspring. Maternally neglected offspring die if not helped, but can regain some of their survival by being helped. Such recovery by being helped has been documented in cooperatively breeding birds (Russell *et al.*, 2007). At the start of the coevolutionary process, the mother is favored to provision all of her offspring. This assumption relies on parental care as an accepted precondition for eusociality (Andersson, 1984).

The interactions in the model are summarized in Fig. 1. Note that maternal influence does not occur through poor provisioning since maternal provisioning is either complete or absent, and thus maternally neglected offspring die if not helped (Fig. 1). Indeed, it will be seen that maternal influence is directed toward first-brood offspring while the mother reduces maternal care toward second-brood offspring.

**Figure 1:**
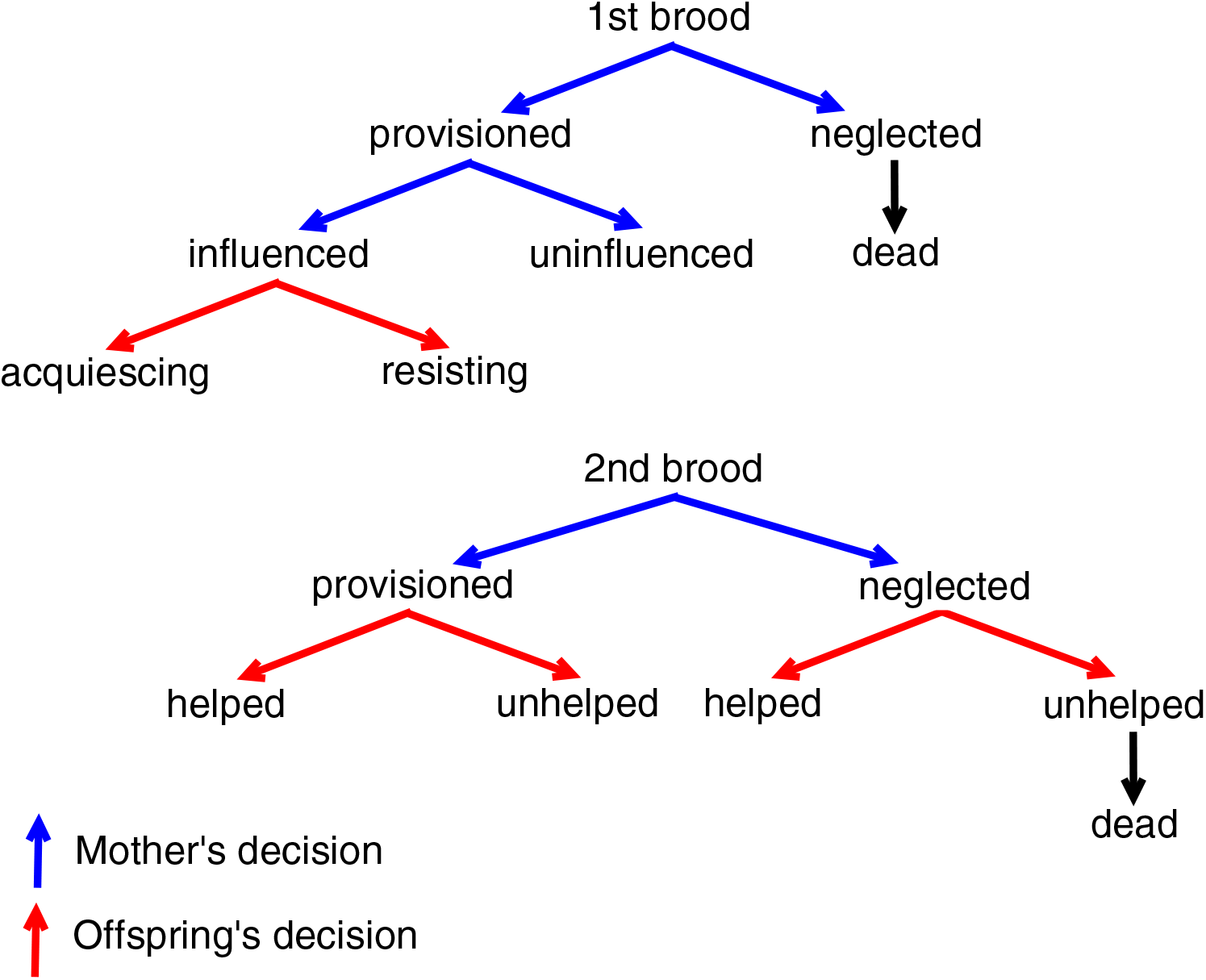
Tree description of the model. See text for details.

The central assumption of the model is the following: I assume that maternally neglected offspring use help more efficiently than maternally provisioned offspring. Consequently, for a unit of food received from helpers, the survival of maternally neglected offspring increases more than that of maternally provisioned offspring. This assumption relies on the expectation that maternally neglected offspring are under stronger pressure to use this food in order to regain survival.

### Maternal influence and offspring resistance

To capture all components of selection on the traits in the model, it is enough to monitor four classes of individuals: (1) young mothers, who produce first-brood offspring; (2) old mothers, who produce second-brood offspring; (3) first-brood subjects (or just subjects), who are the subset of first-brood offspring that can be influenced by the mother (e.g., female offspring as for hymenopteran eusociality, or both female and male offspring as for isopteran eusociality); and (4) second-brood offspring. These four classes are respectively indexed by *i* = m, M, 1, 2.

A focal young mother influences a first-brood subject with probability *p*_m_ to delay dispersal from its natal nest. Here I make use of a notation that I will use throughout: for each trait, the first subscript indicates the class of the individual that controls the trait, while the trait without a class subscript refers to the population average (Table 1). An influenced subject resists with probability *q*_1_ and leaves its natal nest without delay. Alternatively, an influenced subject acquiesces with probability 1 − *q*_1_ and stays in its natal nest for some portion of its adulthood. An acquiescing subject expresses parental care (i.e., provisioning) while in its natal nest with some probability (the evolution of this probability is studied elsewhere; González-Forero, 2014). As stated above, this parental care is directed toward the available brood which are second-brood offspring.

**Table 1:** Notation for the traits.

| In a focal individual | Population average | Definition |
| --- | --- | --- |
| $p_m$ | $p$ | Probability that a mother influences first-brood subjects |
| $q_1$ | $q$ | Probability that an influenced subject resists the influence |
| $a_m$ | $a$ | Fraction of maternal resource allocated to first-brood subjects |
| $e_{m1}$ | $e_1$ | Fraction of the allocated resource to first-brood subjects that the mother spends producing them (she spends the rest provisioning them) |
| $e_{m2}$ | $e_2$ | Fraction of the allocated resource to second-brood offspring that the mother spends producing them |
| $x_1$ | $x$ | Probability that a first-brood subject stays spontaneously |

The survival of a second-brood, maternally provisioned offspring increases by an amount *b*_p_ for each helper that helps it individually, while that of a maternally neglected one increases by an amount *b*_n_. Such *b*_p_ and *b*_n_ specify the benefit from being helped. By the assumption that maternally neglected offspring use help more efficiently than maternally provisioned offspring, I let *b*_n_ > *b*_p_. An increasing number of helpers increases the actual benefit received by helped offspring. Each helper splits uniformly its provisioning effort across second-brood offspring; thus, an increasing number of second-brood offspring decreases the actual benefit received by helped offspring (Charlesworth, 1978). The survival of a helper, which is the probability that the helper becomes a parent itself, decreases by *c*_p_ or *c*_n_ for helping maternally provisioned or maternally neglected offspring respectively. So, *c*_p_ and *c*_n_ define the costs of acquiescence which include the effect of missed reproductive opportunities due to delayed dispersal. Costs of acquiescence that depend on recipient’s condition (*c*_p_ or *c*_n_) allow to account for recipients being more or less demanding of food depending on their condition. Importantly, I assume that maternal influence and offspring resistance are costless (the effect of their costs is explored elsewhere; González-Forero and Gavrilets, 2013, González-Forero, 2014).

### Maternal resource allocation

After recently mated females compete globally for patches, each successful female secures a patch with resources. Of these resources, the female has an amount *R* in energy units to produce and to provision both first-brood subjects and second-brood offspring. The young mother allocates a fraction *a*_m_ of resource *R* to first-brood subjects, and the remaining fraction to the second brood. Of the resource allocated to first-brood subjects, the mother allocates a fraction *e*_m1_ into producing the offspring while she allocates the rest into provisioning them. Similarly, of the resource allocated to the second brood, the mother allocates a fraction *e*_m2_ into producing the offspring and the rest into provisioning them (writing *e*_m2_ instead of *e*_M2_ makes no difference because it is the same mother that controls the trait). The energetic cost of producing an average offspring is *γ*_*π*_ and that of provisioning it is *γ*_p_. For simplicity, I assume that the mother produces a continuous rather than a discrete number of offspring. Hence, the number of offspring of class *i* = 1, 2 produced by the mother are respectively

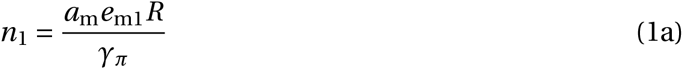

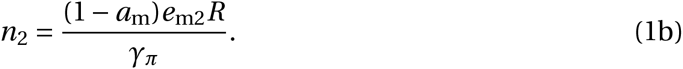

Thus, the total number of monitored offspring produced by a mother is *n* = *n*_1_ + *n*_2_ = (*R*/*γ*_*π*_)[*a*_m_*e*_m1_ + (1 − *a*_m_)*e*_m2_]. The fraction of monitored offspring that are produced as first-brood subjects is *α* = *n*_1_/*n* = *a*_m_*e*_m1_/[*a*_m_*e*_m1_ + (1 − *a*_m_)*e*_m2_]. The number of offspring of class *i* = 1, 2 that the mother provisions herself is

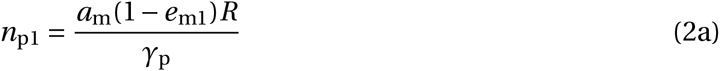

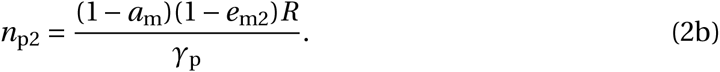

Since the number of maternally provisioned offspring cannot be greater than the number of offspring (*n*_p*i*_ ≤ *n*_*i*_), allocation to offspring production has by definition a lower bound given by *e*_m*i*_ ≥ *γ*_*π*_/(*γ*_*π*_ + *γ*_p_), provided that the mother invests in the two broods (i.e.,0 < *a*_m_ < 1).

In the model, the benefit received by helped offspring (*b*_p_, *b*_n_) and the cost of acquiescence paid by helpers (*c*_p_, *c*_n_) depend on the condition of the helped offspring (i.e., maternally provisioned or maternally neglected). Hence, for a focal helper, the average benefit and cost across its helped recipients depend on maternal resource allocation. Provided that the mother produces the two broods (so 0 < *a*_m_ < 1), the probability that a class-*i* offspring is maternally provisioned is *ζ*_*i*_ = *n*_p*i*_ /*n*_*i*_ = (*γ*_*π*_/*γ*_p_)(1 − *e*_m*i*_)/*e*_m*i*_. Then, for a focal helper, the average cost of acquiescence and the average benefit for its helped recipients are

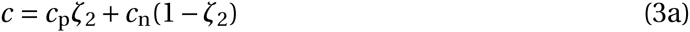

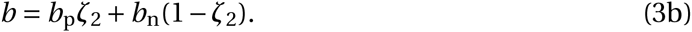

Note that the benefit *b* and cost *c* are under maternal genetic control because they are functions of maternal allocation to offspring production (*e*_m*i*_) and provisioning (1 − *e*_m*i*_).

### Model implementation

I study the coevolution of the population average maternal influence (*p*), offspring costless resistance (*q*), and maternal resource allocation (*a*, *e*_1_, *e*_2_). I assume them to be additive, uncorrelated, quantitative genetic traits. The population is finite, reproduction is sexual and deterministic so genetic drift is ignored, and the genetic system is either diploid or haplodiploid. The total resource in the environment measured in energy units is constant and is divided uniformly among successfully competing mated females, which regulates population growth. I use the approach of Taylor and Frank (1996) to obtain differential equations describing evolutionary change. This approach requires differentiation, so in order to apply it, I use conservative approximations of offspring survival to make it always differentiable. The mathematical details of the model are given in the Appendix. Additional notation is summarized in Table 2.

**Table 2:**
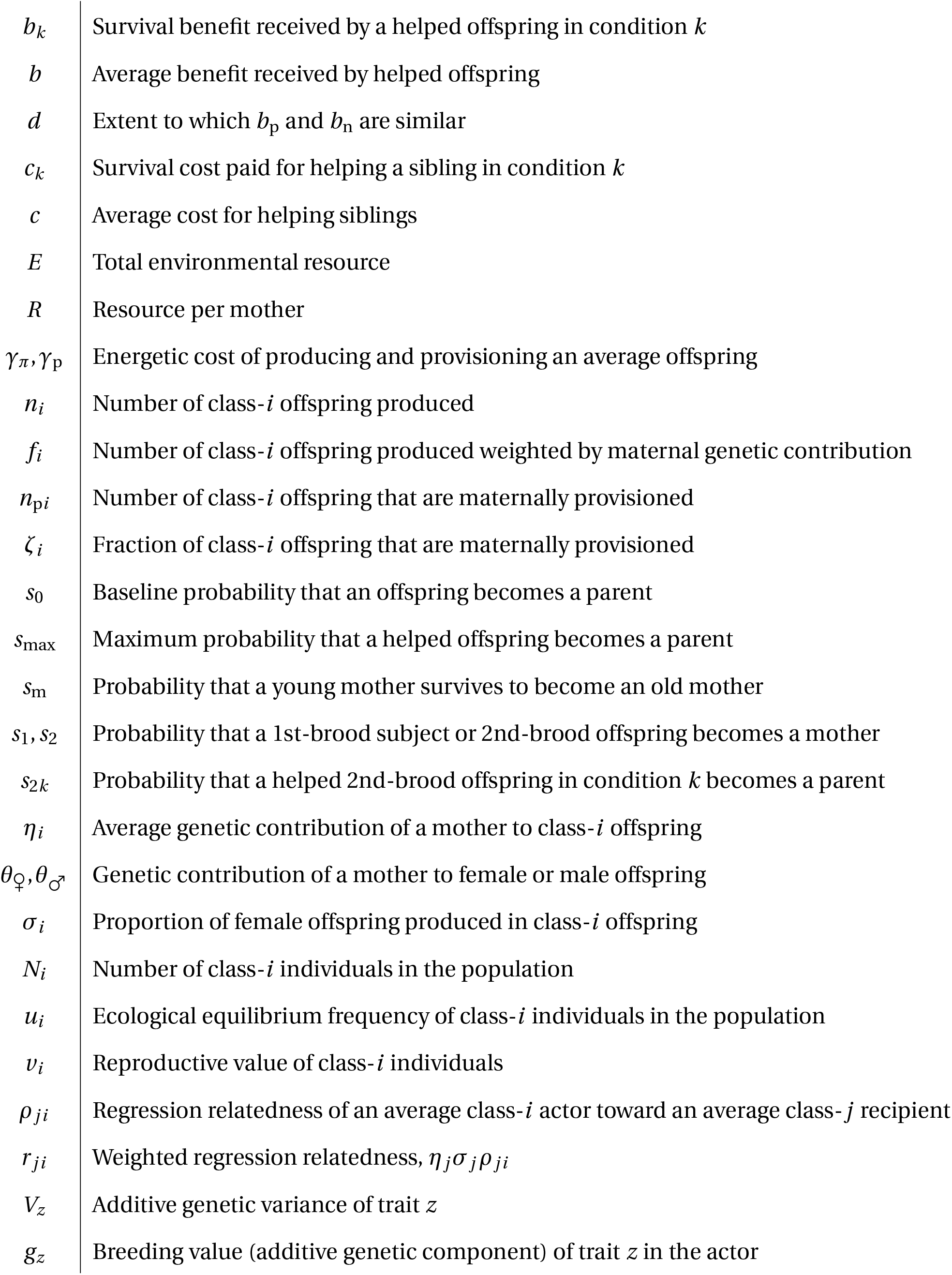
Additional notation. Offspring condition is *k* = p, n if maternally provisioned or maternally neglected.

I solve numerically the differential equations describing evolutionary change. To properly initialize the numerical solutions, I first let maternal resource allocation settle at an equilibrium by allowing it to evolve at a fast pace during 1000 generations without genetic variation for the other traits. Then, I introduce genetic variation for manipulation and resistance. Supporting Figs. referenced below are in the Supporting Information 1 (SI1). The computer code used to generate all figures is in the Supporting Information 2 and 3 (SI2 and SI3).

## Results

The coevolution of maternal influence (*p*), offspring costless resistance (*q*), and maternal resource allocation (*a*, *e*_1_, *e*_2_) yields the following result. At the start of the evolutionary process, both maternal influence and offspring resistance evolve (lines on red shade of Fig. 2a). Hence, there is a mother-offspring conflict over offspring helping behavior (red shade on Fig. 2a-f), and so maternal influence constitutes maternal manipulation during this stage. Manipulation produces a few helpers while resistance is still ineffective (green line on red shade of Fig. 2b). With help available, the mother reduces her maternal care toward second-brood offspring (red line on red shade of Fig. 2c). Thus, first-brood helpers help an increasing proportion of maternally neglected second-brood offspring (*ζ*_2_ decreases from 1). Since by assumption maternally neglected offspring use help more efficiently, the average benefit received by second-brood offspring increases [blue line in Fig. 2d; see eq. (3b)]. The average benefit reaches a sufficiently high level that resistance becomes disfavored [non-shaded area in Fig. 2a; see eq. (A10b)]. Because there are no costs of resistance, resistance being disfavored means that the conflict disappears and maternal influence stops being manipulation as defined above. First-brood subjects become effectively sterile because the cost for helping maternally neglected offspring is here maximal (*c*_n_ = *s*_0_) and so the probability that first-brood subjects become parents (i.e., their survival to parenthood) evolves to zero (Fig. 2e). Daughters that successfully become mothers are no longer raised by the mother but by sterile workers (Fig. 2f). At the end of this coevolutionary process, there is reproductive division of labor where reproductives (i.e., non-sterile offspring, which are the second brood) are produced by the mother but are raised by workers (Fig. 2b,c,e), workers do not reproduce (Fig. 2e), and workers are maternally induced to help but are not selected to resist (Fig. 2a). Because of the final lack of conflict, the final maternal influence fits the notion of signaling: it is a non-conflicting influence that evolved for the purpose of altering offspring’s phenotype while offspring evolved to attend to it (Maynard Smith and Harper, 2003). Therefore, despite costless resistance, maternal manipulation generates stable eusociality and an associated maternal signal that induces offspring to be workers. This process occurs both in haplodiploids and diploids (Supporting Figs. 3-5).

**Figure 2:**
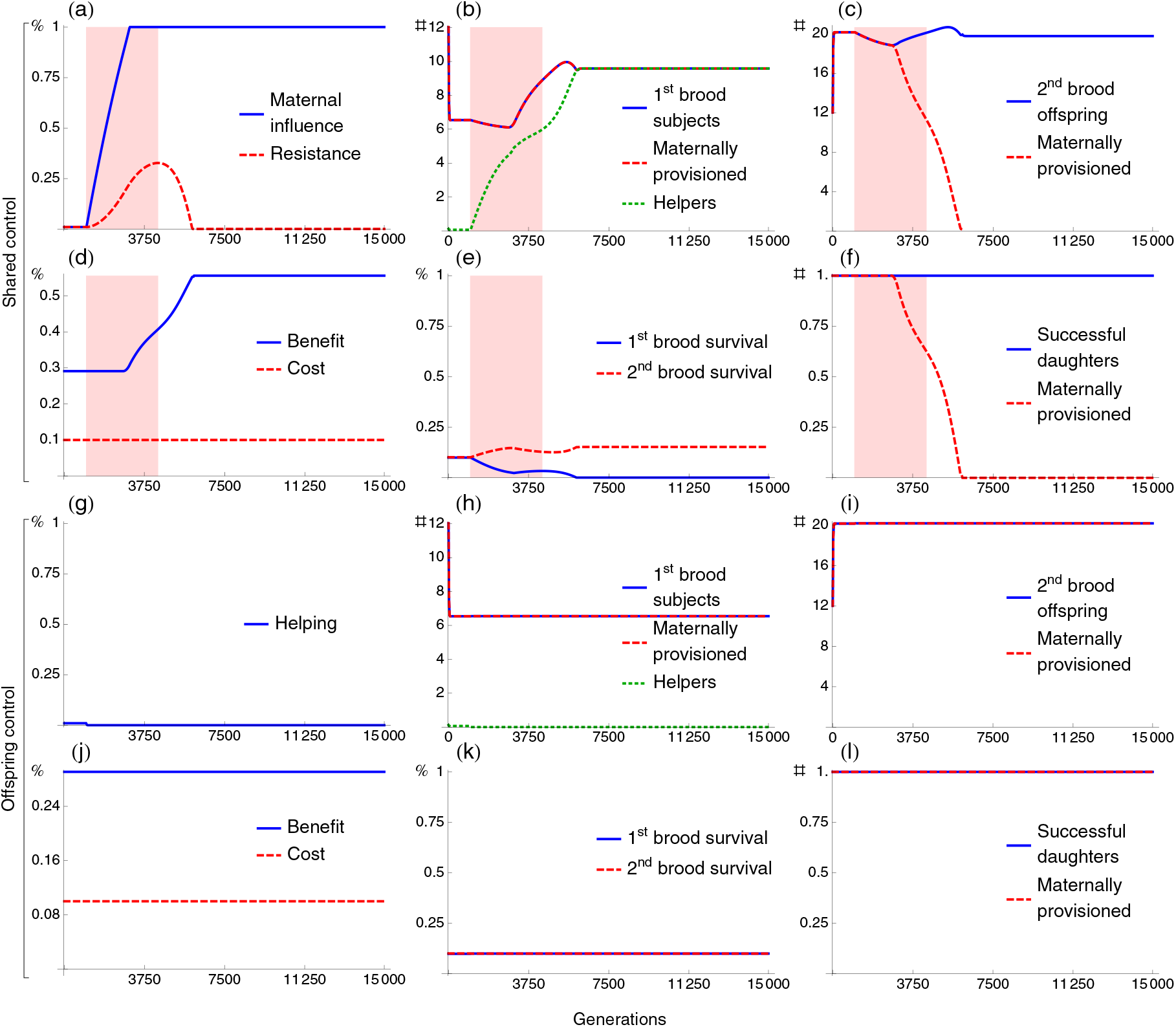
Stable eusociality via maternal manipulation with costless resistance. The plots show population-average values vs. generations. In the two top rows, offspring can be influenced by their mother to stay to help (shared control) (a-f). In the two bottom rows, offspring can stay without being influenced (offspring control) (g-l). In red shades, resistance to the maternal influence is favored to evolve (mother-offspring conflict). Because (a) resistance is initially ineffective, (b) the mother initially has some helpers that allow her to reduce maternal care to the second brood, thereby (d) increasing the benefit that second-brood offspring receive from being helped which (a) eliminates selection for resistance. The genetic system is haplodiploid. Parameter values are in the Supporting Information 1 (SI1).

To assess whether the above process is likely to yield eusociality, I compare the model with two extreme possibilities in which either the mother or the offspring are in full control of offspring’s helping behavior. For the first extreme possibility, I set both the initial resistance and its genetic variation to zero. I refer to this case as maternal control (MC). For the second extreme possibility, I use an otherwise analogous model except that staying in the natal nest is only under offspring control rather than being influenced by the mother (see Offspring control in Appendix). I refer to this case as offspring control (OC). I refer to the intermediate case where maternal influence and offspring resistance can coevolve as shared control (SH). Under the specific parameter values used above for shared control (Fig. 2a-f), eusociality fails to evolve with offspring control (Fig. 2g-l and Supporting Figs. 6,7). Systematic exploration of the parameter space shows that the parameter region in which stable eusociality is obtained is consistently largest with maternal control, followed by shared control, and smallest with offspring control (Fig. 3 and Supporting Figs. 9-14). This result contrasts with previous understanding indicating that the parameter region for stable eusociality should be identical for shared control and offspring control when there are no direct costs associated with resistance (e.g., Craig, 1979, Keller and Nonacs, 1993, Cant, 2006, Uller and Pen, 2011). Specifically, stable eusociality is here obtained with smaller benefit-cost ratios under shared control than under offspring control even though resistance to the maternal influence is entirely costless (note that *b*_p_ and *c*_p_ give the initial benefit and cost for helping because mothers initially provision all their offspring). This occurs more markedly when (1) maternally neglected offspring are substantially more efficient help users than maternally provisioned offspring (i.e., *b*_n_ ≫ *b*_p_), and (2) the survival of maternally provisioned offspring can increase only moderately by being helped (i.e., *s*_0_ → *s*_max_; see Figs. 3a,b and Supporting Figs. 11a,b and 13a,b). The latter condition states that the survival of maternally provisioned offspring must be close to saturation, which occurs when their survival without help (*s*_0_) is already close to the maximum *s*_max_ they can have with help.

**Figure 3:**
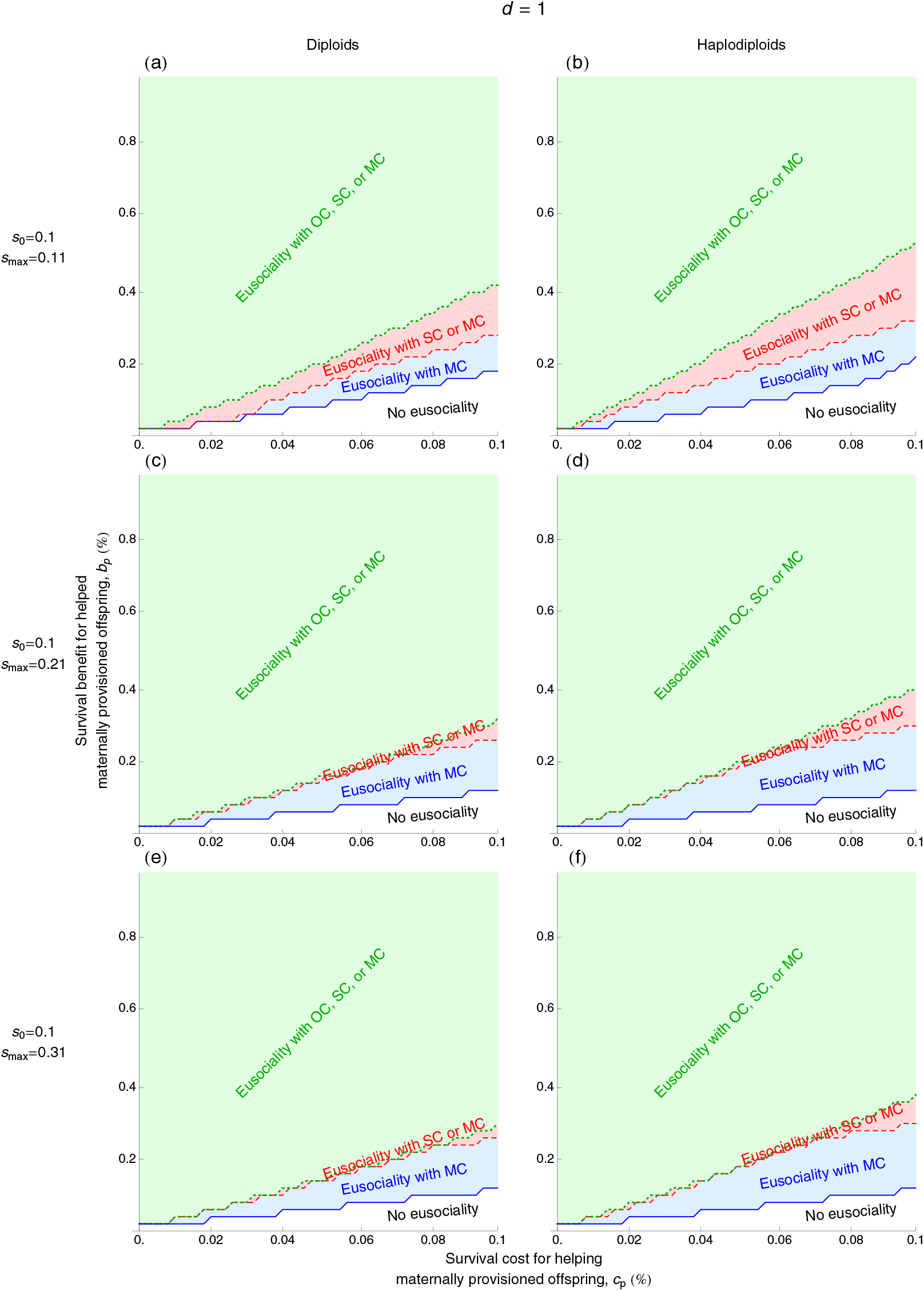
Stable eusociality via maternal manipulation can be obtained under smaller benefit-cost ratios than via offspring control despite costless resistance. The graphs show the outcome across values of the survival benefit for helped maternally provisioned offspring (*b*_p_) vs. the survival cost for helping maternally provisioned offspring (*c*_p_). In blue shade, eusociality is obtained with maternal control of offspring helping behavior (MC). In red shade, eusociality is obtained with either shared control (SC) or maternal control (MC). In green shade, eusociality is obtained with either offspring control (OC), shared control (SC), or maternal control (MC). When the cost for helping maternally provisioned siblings is maximal (here *c*_p_ = *s*_0_ = 0.1), the initial workers are sterile. An evolutionary outcome is here considered eusociality if at the end of the process the two broods are present (*n*_*i*_ ≥ 1) and if there is at least one sterile helper in the first brood [*n*_p1_*p*(1 − *q*) ≥ 1; sterility occurs because in all panels *c*_n_ = *s*_0_ = 0.1]. For the left column, the genetic system is diploid (a,c,e). For the right column, the genetic system is haplodiploid (b,d,f). In all panels, *s*_0_ = 0.1. For the top row, *s*_max_ = 0.11 (a,b), the middle row *s*_max_ = 0.21 (c,d) and the bottom row *s*_max_ = 0.31 (e,f). Finally, *b*_n_ = *b*_*p*_ *d s*_max_/(*d s*_max_ − *s*_0_) and *d* = 1. The remaining parameter values are in the SI1.

## Discussion

In eusocial taxa, queens exert substantial influence on their colonies by prompting offspring to develop or maintain worker phenotypes (e.g., Wilson, 1971, Fletcher and Ross, 1985, O’Donnell, 1998, Le Conte and Hefetz, 2008, Van Oystaeyen *et al.*, 2014). Yet, how queen influence evolved and why it is so common remains poorly understood (Oi *et al.*, 2015). One possible reason for the commonality of maternal influence is that it is a causal factor in the origin of eusociality (Alexander, 1974, Michener and Brothers, 1974, Linksvayer and Wade, 2005, Russell and Lummaa, 2009). Eusociality can evolve under relatively lax conditions if the maternal influence is manipulative and resistance to it is costly (Charlesworth, 1978, González-Forero and Gavrilets, 2013). Otherwise, with costless resistance, eusociality via manipulation is expected to be evolutionarily unstable (Trivers, 1974, Craig, 1979, Keller and Nonacs, 1993). In contrast to this expectation, I show here that maternal manipulation with costless resistance can yield stable eusociality. The reason is that maternal care reduction increases the benefit that offspring receive from help (further explained below). This result relies on the assumption that offspring receiving no maternal care use help more efficiently than offspring receiving maternal care.

### Conflict resolution: from manipulation to honest signaling

Depending on whether helping behavior is controlled by mother, offspring, or both, four broad cases can be considered. First, with *maternal* control and ignoring offspring resistance, eusociality evolves under particularly small benefit-cost ratios (e.g., Charlesworth, 1978, Kapheim *et al.*, 2015; eusociality with MC in Fig. 3). Second, with *offspring* control, eusociality requires larger benefit-cost ratios (e.g., Charlesworth, 1978, Kapheim *et al.*, 2015; eusociality with OC in Fig. 3). Third, with *shared* control between mother and offspring and *costly* resistance, eusociality evolves and is stable under intermediately small benefit-cost ratios (e.g., González-Forero and Gavrilets, 2013, González-Forero, 2014). Fourth, with *shared* control and *costless* resistance, eusociality evolves and is stable under exactly the same benefit-cost ratios as with offspring control (e.g., Craig, 1979, Keller and Nonacs, 1993, Godfray, 1995, Cant, 2006, Uller and Pen, 2011, González-Forero and Gavrilets, 2013). These scenarios have suggested that, when resistance is costless, considering offspring control should be sufficient to explain the evolution of offspring helping behavior (Trivers and Hare, 1976, Craig, 1979, Cant, 2006, Uller and Pen, 2011, Kuijper and Hoyle, 2015). In contrast, the results obtained here show that with shared control and costless resistance, eusociality can still evolve and be stable under intermediately small benefit-cost ratios. Indeed, with maternal manipulation, an initially moderate benefit can evolve and increase sufficiently that helping becomes favored. This is possible because the mother initially produces ineffectively resisting helpers that allow her to reduce maternal care, thereby increasing the benefit and stabilizing eusociality. Without maternal manipulation, a moderate benefit does not increase to favor helping. Since in this case the mother does not have helpers, she does not evolve reduced maternal care that would allow the benefit to increase.

Hence, the evolution of the benefit eliminates the mother-offspring conflict introduced by manipulation. In a previous study where the benefit is controlled by helpers because they control their helping efficiency, the mother-offspring conflict also disappears (González-Forero, 2014). In the present study the benefit is genetically controlled by the mother, since maternal care determines the efficiency of help use by offspring [see eq. (3b)]. These studies fall within a larger class of mathematical models showing that the evolution of fitness payoffs (here *b* and *c*) can reduce, eliminate, or increase the level of conflict (Worden and Levin, 2007, Akçay and Roughgarden, 2011, González-Forero, 2014, Stewart and Plotkin, 2014).

After the mother-offspring conflict disappears, the maternal influence becomes a signal (*sensu* Maynard Smith and Harper, 2003). This signal only informs first-brood offspring that they can have recipients of help if they stay to help. Second-brood offspring do not receive the signal. Helping is then favored as long as it is expressed only when receiving the signal, otherwise it could be expressed by second-brood individuals who have no brood to help. The signal can thus be maintained in evolutionary time to maintain helping (González-Forero, 2014). Given the final absence of mother-offspring conflict over helping behavior, mother and offspring can then evolve in a mutually beneficial way allowing the signal to remain honest. Mutually beneficial coevolution permits subsequent elaborations of the maternal signal. If offspring evolve the ability to provision their mother, offspring could become sensitive to maternal fertility since they affect it directly (see below). Then, the maternal signal could evolve into an honest signal of queen fertility. This pathway links the origin of eusociality to the evidence suggesting that queen pheromones act as honest signaling of the queen’s reproductive health (Heinze and d’Ettorre, 2009, van Zweden *et al.*, 2014).

### Why can eusociality via maternal manipulation be stable when resistance is costless?

The model shows that selection for resistance disappears as the mother reduces maternal care and reallocates resources into producing more offspring. The benefit increases as maternal care decreases because by assumption maternally neglected offspring use help more efficiently than maternally provisioned offspring. The benefit can increase sufficiently that selection for resistance is eliminated because first- and second-brood offspring are siblings (in particular, full siblings for the parameter values explored here) [Hamilton, 1964; see eq. (A10b)]. Given a mathematical equivalence between kin and group selection (Frank, 2012), resistance becomes disfavored once the benefit is large enough that kin or group selection favor acquiescence to the maternal influence. In the model, acquiescence becomes favored because of maternal care reduction but not because of maternal fertility increase. There are two reasons for this. First, maternal fertility remains largely constant because maternal resource decreases with population growth. Maternal resource is obtained from environmental resource divided among mothers so it depends on population size. There is a trade-off between offspring production and provisioning [defined by *e*_m*i*_ in eqs. (1) and (2)], so reduction in provisioning releases resources for offspring production (see Savage *et al.*, 2015 and Kramer *et al.*, 2015). The population grows once the mother starts to reduce care toward second-brood offspring which allows her to produce more offspring (Supporting Fig. 4i). Then, maternal resource becomes smaller with population growth which limits the ability of the mother to increase her fertility. Consequently, the number of first-brood offspring *n*_1_ changes little (Supporting Fig. 4f) as maternal resource *R* decreases with an increasing population size (Supporting Fig. 4n), while the number of maternally provisioned second-brood offspring *n*_2p_ decreases to zero (Supporting Fig. 4h). Therefore, although the benefit *b* can increase as the number of first-brood offspring increases, the observed increase in the benefit *b* is primarily due to maternal care reduction. This effect of competition would not be easily captured by assuming an infinite or constant population size or by imposing a carrying capacity.

Second, acquiescence does not become favored because of an increase of maternal fertility since the benefit *b* that renders resistance disfavored [eq. (A10b)] is not a fertility benefit to the mother and is not weighted by relatedness to the mother. Instead, this benefit *b* is a survival benefit to siblings and is weighted by relatedness to siblings. In the model, helpers do not directly increase maternal fertility. To see this, note that, from eqs. (1), maternal fertility *f*_*i*_ is constant with respect to offspring resistance *q*_1_. Helpers affect maternal fertility only indirectly by allowing the mother to decrease maternal care and redirect the freed resources into additional offspring production. Thus, maternal fertility increase depends on whether the mother chooses to use the help by reducing care and increasing her fertility. Because this choice is here entirely genetically determined, the mother can only increase her fertility as she acquires the genes for this new choice. So, selection is unable to favor acquiescence due to increased maternal fertility if the fertility benefits to the mother occur only generations later. Now, helpers could directly help maternal fertility if they provisioned the mother thus giving her additional resource for offspring production (e.g., if maternal resource *R* were a function of offspring resistance *q*_1_). However, provisioning the mother could demand a greater effect of the maternal influence than just causing offspring to stay as adults. This is because helpers would have to provision an adult rather than a young which may require additional changes to the normal behavioral repertoire of the offspring (Hunt, 2007). Nevertheless, an important extension of the model is to allow for the evolution of offspring provisioning of the mother as this is a widespread behavior in extant eusocial taxa (Wcislo and Gonzalez, 2006). Such an extension could allow for a marked increase in maternal fertility, which is not recovered in the model (Fig. 2b,c and Supporting Fig. 4f). These observations highlight the importance of detailing how helping occurs and so who the direct recipient of help is: here, it is second-brood offspring rather than the mother.

### The assumption of efficient help use

The assumption that maternally neglected offspring use help more efficiently than maternally provisioned offspring relies on the expectation that maternally neglected offspring are under stronger pressure to regain survival. This assumption must be tested by assessing whether the survival of maternally neglected offspring increases faster than that of maternally provisioned offspring with respect to the ratio of helpers to recipients when this ratio approaches zero (see Supporting Figs. 1 and 2).

The more efficient help use by maternally neglected offspring is a biological assumption that must be tested rather than a mathematical consequence of the model. A similar mathematical consequence of the model is that the marginal benefit received by maternally neglected offspring is larger than that obtained by maternally provisioned offspring. This is because maternally neglected offspring die if not helped and they can reach the same maximum survival of maternally provisioned offspring. Then, it can be checked that, for the differentiable approximations of survival used, the marginal benefit for maternally neglected offspring (which is the negative of the derivative of *s*_2_ with respect to *Q* setting *ζ*_2_ = 0) is larger than that for maternally provisioned offspring even if *b*_n_ = *b*_p_. However, such larger marginal benefit is not enough to eliminate the mother-offspring conflict if *b*_n_ = *b*_p_ (results not shown). Instead, the biological efficiency of help use must be larger for maternally neglected offspring (*b*_n_ > *b*_p_), which can be tested as described in the previous paragraph.

### Model predictions

When the assumption of efficient help use holds, the model makes predictions to discern whether eusociality is likely to have originated from maternal manipulation rather than from offspring control, particularly when resistance is costless. One prediction is that stable eusociality via manipulation and maternal care reduction is more likely when the survival of maternally provisioned offspring can increase only moderately with help; that is, their survival must be close to saturation (Fig. 3a,b and Supporting Figs. 11a,b and 13a,b). On the contrary, eusociality via offspring control does not require that the survival of maternally provisioned offspring is close to saturation (Fig. 3 and Supporting Figs. 9-14).

In addition, the disappearance of the mother-offspring conflict predicts the occurrence of “conflict relics”. By this I mean a trait (e.g., morphological, molecular, or behavioral) that ancestrally served as an adaptation for manipulation or resistance but lost this function. For example, conflict relics predict the putative conflicting genes to have a high within species genetic diversity (reflecting conflict) that is shared between recently diverged species (reflecting that conflict is ancestral but not current) (see Ostrowski *et al.*, 2015). Because conflict relics are not expected if eusociality originates via offspring control, conflict relics also allow to disentangle manipulation and offspring control as a source of eusociality, even with costless resistance.

### Further biological implications

Several points in the model warrant further comment. First, reproductive value does not drive the process described here although it evolves and becomes small for helpers and large for recipients. Previous theory shows that if helping entails *fertility* costs and benefits, helping is favored when helpers’ reproductive value is lower than that of helped individuals (West Eberhard, 1975, Frank, 1998), which has prompted hypotheses for the evolution of eusociality (e.g., Holman, 2014). Here helping entails only *survival* costs and benefits, and so reproductive values cannot change the direction of selection and instead the class equilibrium frequencies (*u*_*i*_) play the analogous role [i.e., the derivatives of *f*_*i*_ in eqs. (A9) are here zero]. Still, class equilibrium frequencies do not cause the observed change in selection for resistance since here they affect the direction of selection via the sex ratio in the two broods [i.e., the *η*_*j*_ *σ*_*j*_ occurring in *r*_*j,i*_ in eqs. (A10)], which I assumed even and constant. Yet, in the model, first-brood individuals evolve low reproductive values as their survival decreases, while second-brood individuals evolve high reproductive values as their survival increases [eqs. (A16c) and (A16d) and Supporting Figs. 2l and 3l], which matches the expected pattern.

Second, the model considers a finite population, the size of which is regulated by the finite environmental resource without imposing a carrying capacity. Then, population size and the number of individuals of different classes can evolve as trait values change (Supporting Figs. 2i,j and 3i,j). This aspect differs from previous models that usually assume infinite or constant population sizes. Third, genetic variances are important on whether eusociality is stabilized. Although the model’s complexity prevents analytical treatment, a simpler version of the model suggests that stable eusociality via manipulation and care reduction requires a condition of the form *br* + (1 − *q*_0_)*A* > *c* which allows acquiescence to become favored as the benefit evolves (see eq. A3.50e in González-Forero, 2013). In this inequality, *r* is relatedness of first- to second-brood offspring, *q*_0_ is the initial resistance, and *A* is proportional to the ratio of genetic variances of maternally controlled traits over the genetic variance of offspring resistance. This inequality suggests that large genetic variances for maternally controlled traits relative to offspring controlled traits would favor the disappearance of conflict via this process. Fourth, the model describes parental care as provisioning, but it can be equivalently taken as nest defense directed to individuals (Cocroft, 2002). Parental care in the form of defense is important because it is thought to have been key for the origin of isopteran eusociality (Korb *et al.*, 2012). In this interpretation of the model, reduced maternal care toward second-brood offspring refers to reduced maternal investment into defending individual second-brood offspring.

Finally, two underlying assumptions of the models can be relevant to account for the high incidence of eusociality in hymenoptera and its occurrence in termites. Without maternal influence, a mutant gene for helping must have a dual function: to trigger the expression of help and to be expressed only in first-brood individuals of the right sex. This expression pattern can occur if first-brood individuals of the right sex in the ancestral population already use environmental cues that properly trigger the helping gene expression. With maternal influence, it is the maternal influence gene that must have the analogous dual function: to trigger the expression of maternal influence and to be expressed so that only first-brood offspring of the right sex are influenced. This dual function of a maternal influence gene is particularly feasible in hymenoptera. Indeed, hymenopteran mothers control the sex of their offspring by fertilizing eggs (Verhulst *et al.*, 2010) and their first offspring are often female (Hunt, 2007). Hence, the dual function for the maternal influence gene occurs if the gene is expressed only early in the reproductive phase of a hymenopteran mother. In diploids, the dual ability of the maternal influence gene can also be facilitated by early expression if the early brood is composed by the sex or sexes providing parental care. This requirement is likely to have been met by isopteran ancestors given their probable ancestral biparental care (Klass *et al.*, 2008).

## Conclusion

If maternally neglected offspring are particularly efficient help users, maternal manipulation and maternal care reduction can generate stable eusociality when resistance to manipulation is costless. This scenario requires ancestral parental care, and that maternal manipulation can be executed and favored. With these conditions, ancestral manipulation can then evolve into honest maternal signaling.

## Acknowledgments

I thank Sergey Gavrilets, Kelly Rooker, Jeremy Auerbach, Xavier Thibert-Plante, Laurent Lehmann, Charles Mullon, Simon Powers, Slimane Dridi, Matthias Wubs, Valeria Montano, Danielle Mersch, Laurent Keller, Luke Holman, and two anonymous reviewers for discussion or comments on the manuscript that vastly improved it. I thank L. Holman who refereeing the manuscript suggested to keep the term manipulation for conflict situations to better agree with common usage of the concept. I thank S. Gavrilets for access to the Volos computer cluster where I ran numerical simulations conducive to the results reported here. I was funded by a Graduate Research Assistantship from the National Institute for Mathematical and Biological Synthesis (NIMBioS). NIMBioS is sponsored by the National Science Foundation, the U.S. Department of Homeland Security, and the U.S. Department of Agriculture through NSF Award # EF-0832858, with additional support from The University of Tennessee, Knoxville. Volos is funded by the NIH grant # GM-56693 to S. Gavrilets.

## Appendix

### Life history implementation

I separate time into ecological and evolutionary scales. Individuals reproduce in an ecological time scale, and traits change in an evolutionary time scale. I assume that the ecological time scale is much faster than the evolutionary one. Ecological time is discrete, while evolutionary time is continuous. At each ecological time, I monitor the defined four classes of individuals: young mothers, old mothers, first-brood subjects, and second-brood offspring (indexed by *i* = m, M, 1, 2). A mother produces *n*_*i*_ offspring of class *i* (= 1, 2). A fraction *σ*_*i*_ of *n*_*i*_ is female. The average genetic contribution of the mother to class-*i* offspring is *η*_*i*_ [= *σ*_*i*_ *θ*_*♀*_ + (1 − *σ*_*i*_)*θ*♂, where *θ*_*l*_ is the genetic contribution of a mother to sex-*l* offspring; for diploids, *θ*_*l*_ = 1/2, and for haplodiploids, *θ*_♀_ = 1/2 while *θ*♂ = 1]. Maternal fertility through class-*i* offspring is *f*_*i*_ = *η*_*i*_ *n*_*i*_ (Taylor, 1990). Survival of class-*i* offspring (*i* = 1, 2), defined as the probability that a class-*i* offspring becomes a young mother, is *s*_*i*_. The probability that a young mother becomes an old mother is *s*_m_. The number of class-*i* individuals in the population at ecological time *τ* is *N*_*i*_ (*τ*). With **N** = (*N*_m_, *N*_M_, *N*_1_, *N*_2_)^*T*^, then **N**(*τ* + 1) = **WN**(*τ*) where

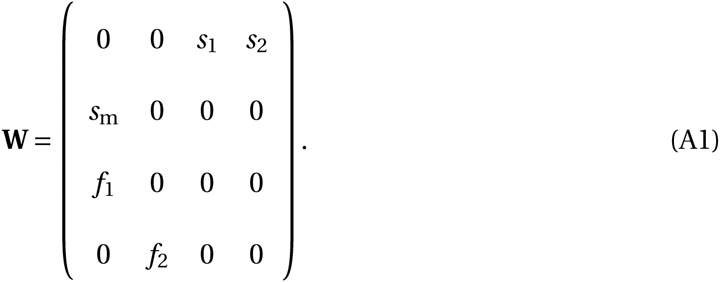

### Survival

I assume that maternal survival *s*_m_ only depends on a constant environmental mortality, and so *s*_m_ is independent of the evolving traits. The probability that a maternally provisioned offspring becomes a parent in the absence of maternal influence or help is *s*_0_ (baseline survival). Since survival *s*_*i*_ (*i* = 1, 2) is the probability of becoming a young mother, the survival of a first-brood subject (who is a female with probability *σ*_1_) is

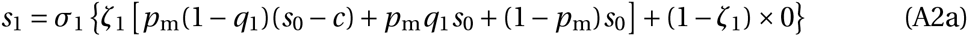

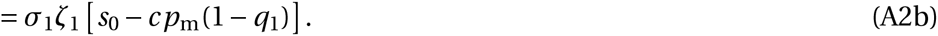

The probability that a second-brood offspring in condition *k* (*k* = p, n) becomes a parent after being helped is *s*_2*k*_. The average resistance probability among the first-brood subjects of a mother is *Q*. So, *p*_m_(1 − *Q*) is the probability that first-brood subjects are helpers. Then, the survival of a second-brood offspring is

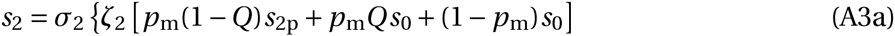

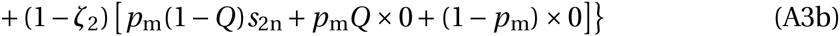

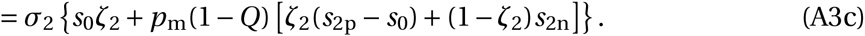

To fully specify the survival of second-brood offspring (*s*_2_), it remains to specify the survival of helped second-brood offspring in condition *k* (*s*_2*k*_).

Let *s*_max_ be the maximum probability of becoming a parent after receiving help (maximum survival). Following Charlesworth (1978), the survival of maternally provisioned offspring after being helped is

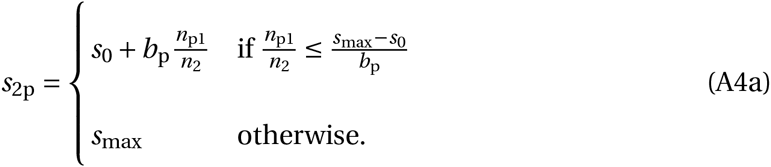

The factor *n*_p1_/*n*_2_ is the number of possible helpers over the number of recipients but since *s*_2p_ is already conditioned on the fact that the second-brood individual is helped, then *n*_p1_ in eq. (A4) gives the number of actual helpers. Survival *s*_2p_ saturates to *s*_max_ if the ratio of helpers to recipients *n*_p1_/*n*_2_ is sufficiently large. The survival of maternally neglected offspring after being helped is

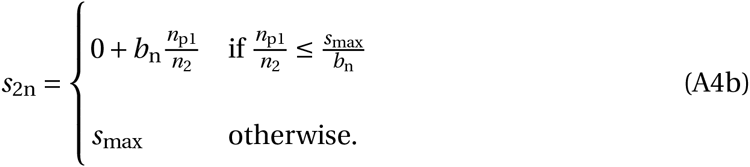

When the ratio of helpers to recipients is sufficiently small [*n*_p1_/*n*_2_ ≤ (*s*_max_ − *s*_0_)/*b*_p_, *s*_max_/*b*_n_], then the survival of a second-brood offspring reduces to

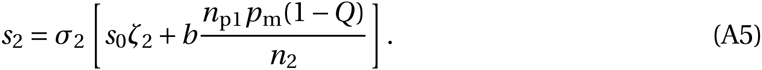

### Survival approximation

Survivals after being helped (*s*_2*k*_) are not differentiable at their switching points when *n*_p1_/*n*_2_ becomes too large. The method of Taylor and Frank (1996) requires differentiation, so I approximate *s*_2*k*_ by always differentiable functions as follows. Denoting *ξ* = *n*_p1_/*n*_2_, *s*_2p_ can be written as a function *s*_2p_(*ξ*) which can be approximated from below by a function of the form

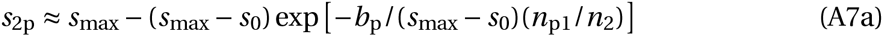

for some *A*_1_, *A*_2_, *A*_3_. Setting *F* (0) = *s*_0_ and *F* (∞) = *s*_max_ yields *A*_1_ = *s*_max_ − *s*_0_ and *A*_2_ = *s*_max_/*A*_1_. Choosing *F*′ (0) = *b*_p_ produces *A*_3_ = *b*_p_/*A*_1_. Proceeding similarly with *s*_2n_ yields the approximations

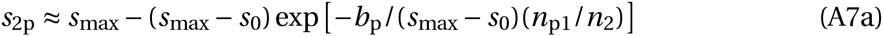

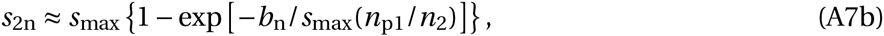

which hold for any *n*_p1_/*n*_2_ > 0 (see Supporting Fig. 2).

### Population regulation

Young mothers compete globally for resources to produce and provision first-brood subjects and second-brood offspring. The environment has a constant amount *E* of resources in energy units that females use for these purposes. Environmental resource *E* is divided uniformly among young mothers, so each young mother has an amount of resource *R* = *E* /*N*_m_. I assume that the population reaches zero growth during ecological time, which occurs when the leading eigenvalue of **W** is one; that is, when *f*_1_*s*_1_ + *s*_m_ *f*_2_*s*_2_ = 1 evaluated at population average values, which is a version of the Euler-Lotka equation (Charlesworth, 1994). Solving for *N*_m_ yields the ecologically stationary number of young mothers

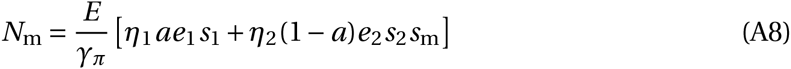

evaluated at population averages. Population size is *N* = *N*_m_ + *N*_M_ + *N*_1_ + *N*_2_, where from **N** = **WN** we have that *N*_M_ = *s*_m_*N*_m_, *N*_1_ = *f*_1_*N*_m_, and *N*_2_ = *f*_2_*N*_M_. Notice that although population size remains constant in ecological time scales, it can evolve in evolutionary time scales as trait values change. From eqs. (1), it follows that the ecologically stationary number of offspring is *n* = 1/[ *η*_1_*αs*_1_ + *η*_2_(1 − *α*)*s*_2_*s*_m_].

### Dynamic equations

I study the coevolution of maternal influence, resistance, and maternal resource allocation (i.e., *p*, *q*, *a*, *e*_1_, and *e*_2_, which denote population averages). As previously stated, I assume they are additive, uncorrelated, quantitative genetic traits. The additive genetic variance of trait *z* is *V*_*z*_ (*z* = *p*, *q*, *a*, *e*_1_, *e*_2_). From the previous section, *R* is a function of population average trait values and is then constant with respect to the actor’s breeding value (i.e., the additive genetic component of the trait in the individual controlling the trait). The equilibrium frequency of class-*i* individuals during the ecological time scale, or simply the class-*i* ecological equilibrium frequency, is *u*_*i*_. The individual reproductive value of class-*i* individuals is *v*_*i*_. *u*_*i*_ and *v*_*i*_ are respectively the right and left eigenvectors of **W** after normalization so that ∑ *u*_*i*_ = ∑ *u*_*i*_*v*_*i*_ = 1 (Leslie, 1948, Taylor, 1990). I assume that mutation and selection are weak. Thus, for evolutionary time *t*, the change in the population average value of trait *z* can be approximated (Taylor and Frank, 1996, Frank, 1997) by

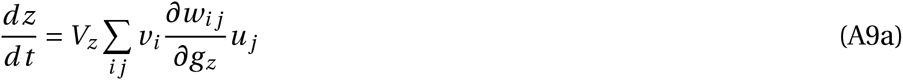

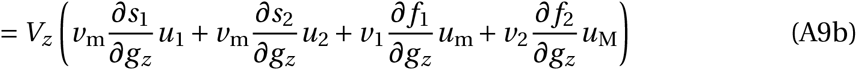

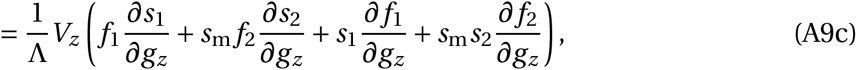

evaluated at population averages, where *w*_*i,j*_ is the *i j* -th entry of **W**, *g*_*z*_ is the actor’s breeding value for *z*, and Λ = 2 + *s*_m_ *f*_2_*s*_2_ is a scaling factor due to population growth. The values of *u*_*i*_ and *v*_*i*_ are found below in Demographic variables.

I solve system (A9) numerically making use of the approximations of *s*_2*k*_ in eqs. (A7) [see Supporting Information 3 (SI3) for computer code]. However, the exact *s*_2*k*_ yield a system that is conceptually useful. Specifically, for *n*_p1_/*n*_2_ ≤ (*s*_max_ − *s*_0_)/*b*_p_, *s*_max_/*b*_n_, using the exact *s*_2*k*_ yields

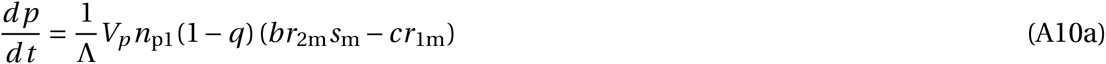

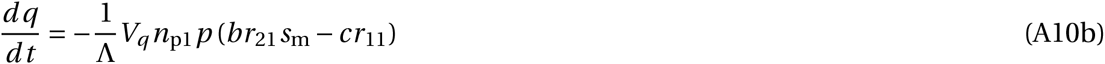

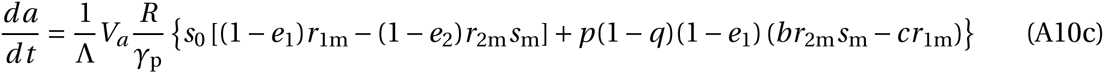

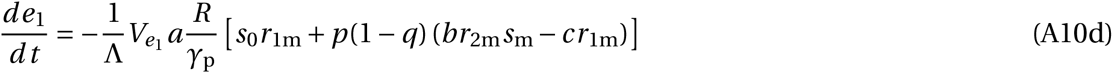

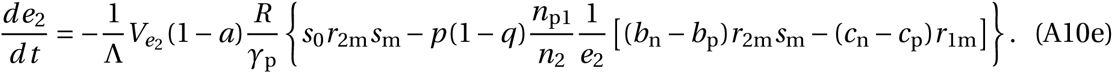

where *r*_*j,i*_ = *η*_*j*_ *σ*_*j*_ *σ*_*j,i*_, *σ*_*j,i*_ = *dz*_*j*_ /*d g_z_i* is the regression relatedness of class-*i* actor to class− *j* recipient, *z*_*j*_ is the trait in the recipient, and *g*_*z*_i is the breeding value in the actor (see SI2 for check of the derivation).

### No helping

By removing maternal influence (setting *p* = 0 and *V*_*p*_ = 0), system (A10) reduces to

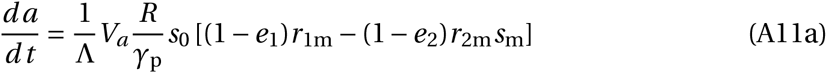

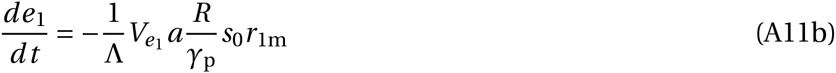

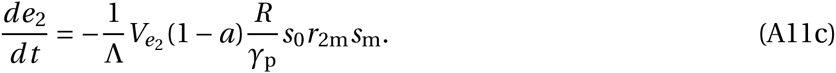

This system evolves to minimal investment in offspring production [i.e., 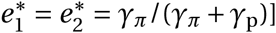 and to either the loss of one brood or to a constant investment in each brood [i.e., *a*^*^ = 0, 1, *a*(0)] depending on how related the mother is to the broods (i.e., depending on whether *r*_1m_ < *r*_2m_*s*_m_, *r*_1m_ > *r*_2m_*s*_m_, or *r*_1m_ = *r*_2m_*s*_m_, respectively). I assume that maternal survival is such that the mother is favored to produce two broods in the absence of helping; so I let *s*_m_ = *r*_1m_/*r*_2m_. For diploids, this means that *s*_m_ = 1 while for haplodiploids *s*_m_ can be smaller than one. A survival *s*_m_ = 1 can refer to the case in which the mother produces and provisions the offspring of both broods at once (mass provisioning), while second-brood offspring hatch from their eggs later. The assumption of *s*_m_ = *r*_1m_/*r*_2m_ can be relaxed in more complex models incorporating selection pressures for producing two broods.

### Offspring control

I consider a modified model where first-brood subjects stay spontaneously (i.e., without maternal influence) in the natal nest for some period of their adulthood. Subjects are here understood as a subset of first-brood offspring in which the staying propensity is expressed (e.g., females only or both sexes). A first-brood subject stays spontaneously with probability *x*_1_. The survival of a first-brood subject is now

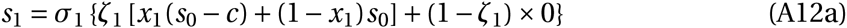

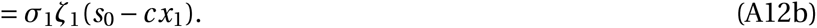

The average probability of staying spontaneously among the first-brood subjects of a mother is *X*. The survival of a second-brood offspring is now

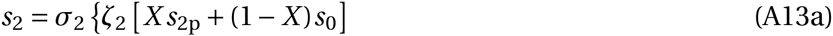

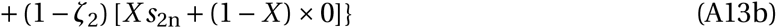

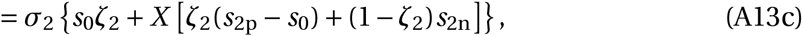

with the exact and approximated *s*_2*k*_ defined as before.

I also solve system (A9) numerically for this model using the approximations of *s*_2*k*_ in eqs. (A7). However, for a sufficiently small ratio of helpers to recipients [*n*_p1_/*n*_2_ ≤ (*s*_max_ − *s*_0_)/*b*_p_, *s*_max_/*b*_n_], using the exact *s*_2*k*_ and letting *x* denote the population average staying probability, the dynamic equations are

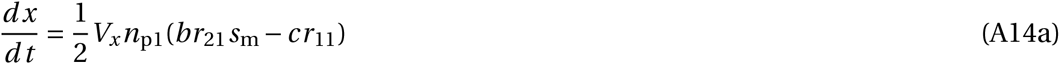

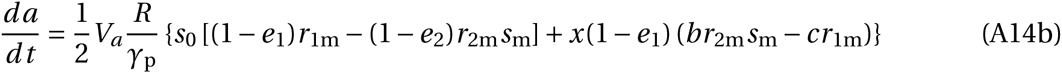

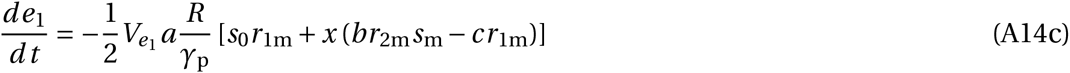

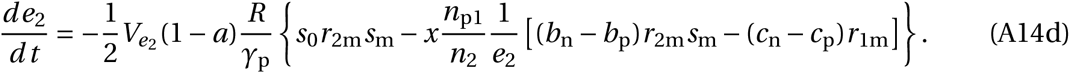

### Demographic variables

The ecologically asymptotic population growth rate is *λ*, which is given by the only real solution of the characteristic equation of **W**; that is, by *λ*^3^ = *λf*_1_*s*_1_ + *s*_m_ *f*_2_*s*_2_. Setting *λ* = 1, the ecological equilibrium frequencies of class-*i* individuals are

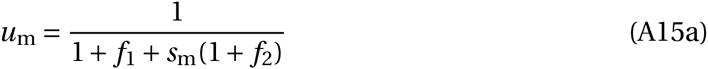

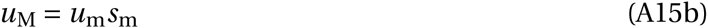

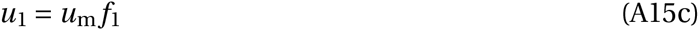

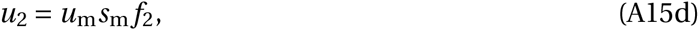

and the reproductive values of class-*i* individuals are

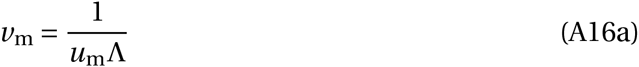

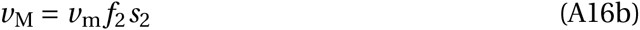

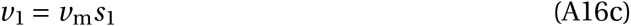

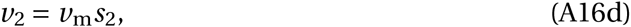

all evaluated at population-average values.

## 1 Parameter values

To calculate regression relatednesses, I use the following expressions:

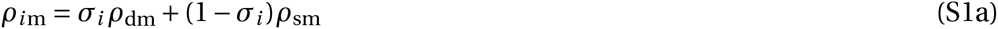

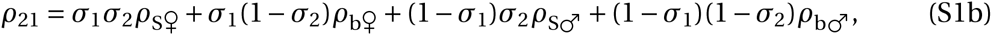

where the subscripts d, s, S, and b refer to daughter, son, sister, and brother respectively. Eqs. (S1) are in terms of standard regression relatedness values that can be obtained from pedigrees given the model assumptions (Hamilton, 1972).

I make the following assumptions. The mother is singly mated. For diploids, both broods have an even sex ratio. For haplodiploids, the second brood has an even sex ratio while the mother directs her influence only to first-brood females (so *σ*_1_ *=* 1). Survival of young mothers to old mothers is such that mothers are initially favored to produce two broods (so *s*_m_ *= r*_1m_/*r*_2m_). However, this value was obtained for the exact survivals, so it is an approximation when using the approximated survival in eqs. (A7) in the main text. Therefore, I let maternal resource allocation evolve alone for 1000 generations to properly initialize the numerical solutions. I let all traits have the same genetic variance to avoid giving an evolutionary advantage to any of them. I let the cost of acquiescence when raising maternally neglected offspring equal the baseline survival (*c*_n_ *= s*_0_), which amounts to saying that helpers of maternally neglected offspring are sterile. I take the initial probability of maternal influence and resistance to be small. I let the initial maternal allocation to be such that the mother produces two equally large broods that she feeds entirely. For simplicity, I let the energetic cost of producing and feeding offspring be the equal. I take the environmental resource to be such that population size is in the tens of thousands.

Finally, I assume that maternally neglected offspring use help more efficiently than maternally provisioned offspring (*b*_n_ *> b*_p_). To reduce the parameter space, I consider two cases: strong and weak advantage in help use efficiency. Specifically, I take *b*_n_ to be as il-lustrated in Supporting Fig. 1. So, the benefit to maternally neglected offspring is *b*_n_ *= b*_p_*d s*_max_/(*d s*_max_ *− s*_0_), where *d =* 1, 2 for strong and weak advantage in help use efficiency respectively.

The remaining parameters are *s*_0_, *s*_max_, *c*_p_, and *b*_p_. From their definitions, they can take values while satisfying 0 *< s*_0_ *< s*_max_ *≤* 1, *c*_p_ *≤ s*_0_, and *b*_p_ *>* 0. With these assumptions, parameter values are those in Supporting Table 1 except when noted otherwise.

**Supporting Table 1:** For Fig. 3 and Supporting Figs. 9-14, *t*_final_ *=* 50 000 while *b*_p_ *∈* [0, 1] and *c*_p_ *∈* [0, *s*_0_]. To properly initialize the numerical solutions, genetic variances are 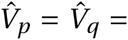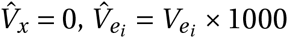, and 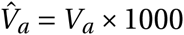 for *t <* 1000. *\**The variance of *e*_*i*_ is scaled so that the additive effect of genes for traits *e*_*i*_ is equal to those for the other traits. †Values taken from Bulmer (1994) following Hamilton (1972).

| $E$ | 100 000 | For diploids | | | |
| --- | --- | --- | --- | --- | --- |
| $V_p, V_q, V_a$ | 0.01 | $\sigma_1, \sigma_2$ | | 0.5 | |
| $V_{e_1}, V_{e_2}^*$ | $0.01\left(1 - \frac{\gamma_\pi}{\gamma_\pi + \gamma_p}\right) = 0.005$ | $\eta_1, \eta_2$ | | 0.5 | |
| $\gamma_\pi, \gamma_p$ | 1 | $\rho_{1m}, \rho_{2m}$ | | 0.5† | |
| $s_0$ | 0.1 | $\rho_{21}$ | | 0.5† | |
| $s_{\max}$ | 0.21 | $s_m$ | | $\frac{r_{1m}}{r_{2m}} = 1$ | |
| $c_p$ | $s_0 = 0.1$ | $b_p$ | | 0.253 | |
| $c_n$ | $s_0 = 0.1$ | $b_n$ | | $b_p \frac{s_{\max}}{s_{\max} - s_0} = 0.483$ | |
| $p(0), q(0)$ | 0.01 | For haplodiploids | | | |
| $e_1(0), e_2(0)$ | $\frac{\gamma_\pi}{\gamma_\pi + \gamma_p} = 0.5$ | $\sigma_1$ | 1 | $\sigma_2$ | 0.5 |
| $a(0)$ | 0.5 | $\eta_{\text{♀}}$ | 0.5 | $\eta_{\text{♂}}$ | 1 |
| $t_{\text{final}} =$ | 15 000 | $\eta_1$ | 0.5 | $\eta_2$ | 0.75 |
| | | $\rho_{\text{dm}}$ | 0.5† | $\rho_{\text{sm}}$ | 1† |
| | | $\rho_{\text{S♀}}$ | 0.75† | $\rho_{\text{b♀}}$ | 0.5† |
| | | $\rho_{\text{S♂}}$ | 0.25† | $\rho_{\text{b♂}}$ | 0.5† |
| | | $\rho_{1m}$ | 0.5 | $\rho_{2m}$ | 0.75 |
| | | $\rho_{21}$ | | 0.625 | |
| | | $s_m$ | | $\frac{r_{1m}}{r_{2m}} \approx 0.8889$ | |
| | | $b_p$ | | 0.291 | |
| | | $b_n$ | | $b_p \frac{s_{\max}}{s_{\max} - s_0} = 0.555$ | |

## 2 Supporting figures

**Supporting Figure 1:**
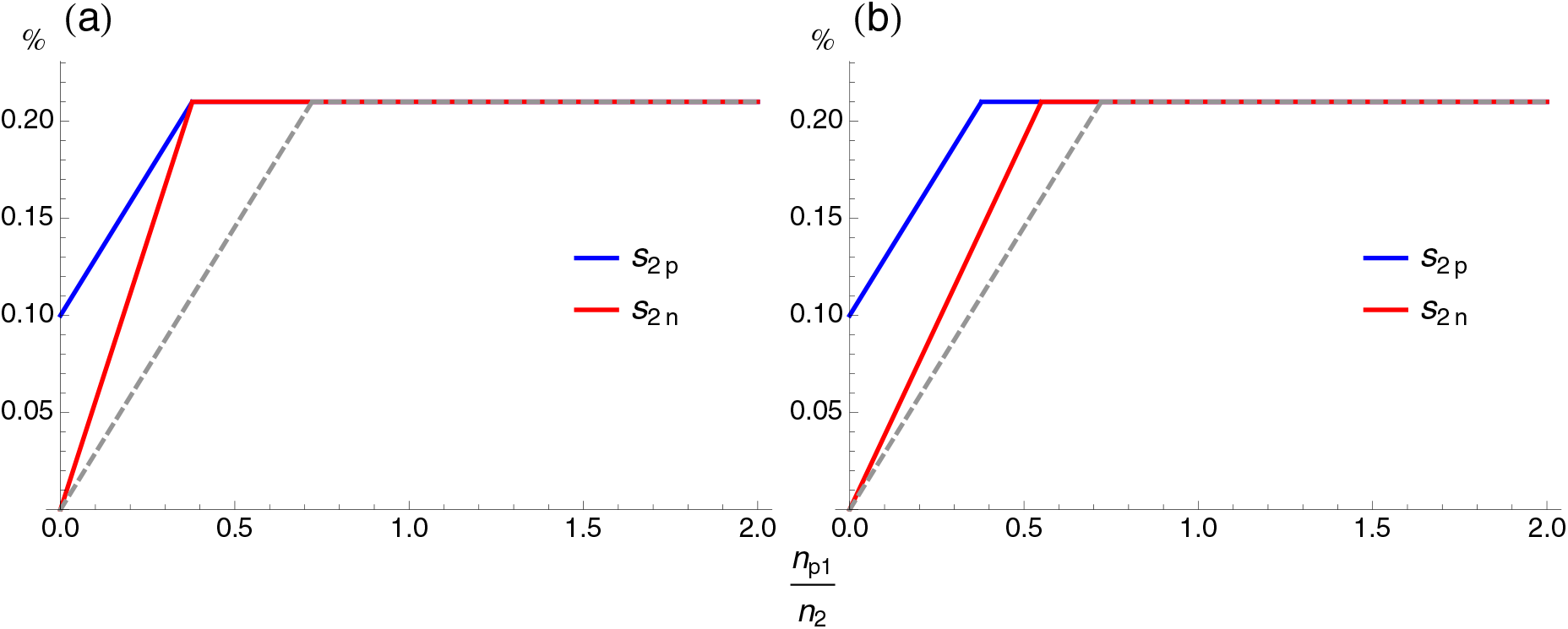
Survival of recipients of help. Plots are the survival of helped second-brood offspring that are maternally provisioned (blue lines) or maternally neglected (red lines) vs. the number of helpers over recipients. The slope of the red line is the survival benefit from being helped for maternally neglected offspring [which for small *n*_p1_/*n*_2_ is *b*_n_ *= b*_p_*d s*_max_/(*d s*_max_ *− s*_0_)]. The advantage in help use efficiency by maternally neglected offspring is either (a) strong (*d =* 1) or (b) weak (*d =* 2). The dashed gray line is the survival of helped maternally neglected second-brood offspring when they have no advantage in help use efficiency (*b*_n_ *= b*_p_). Parameter values are those for haplodiploids in the Supporting Table. 1.

**Supporting Figure 2:**
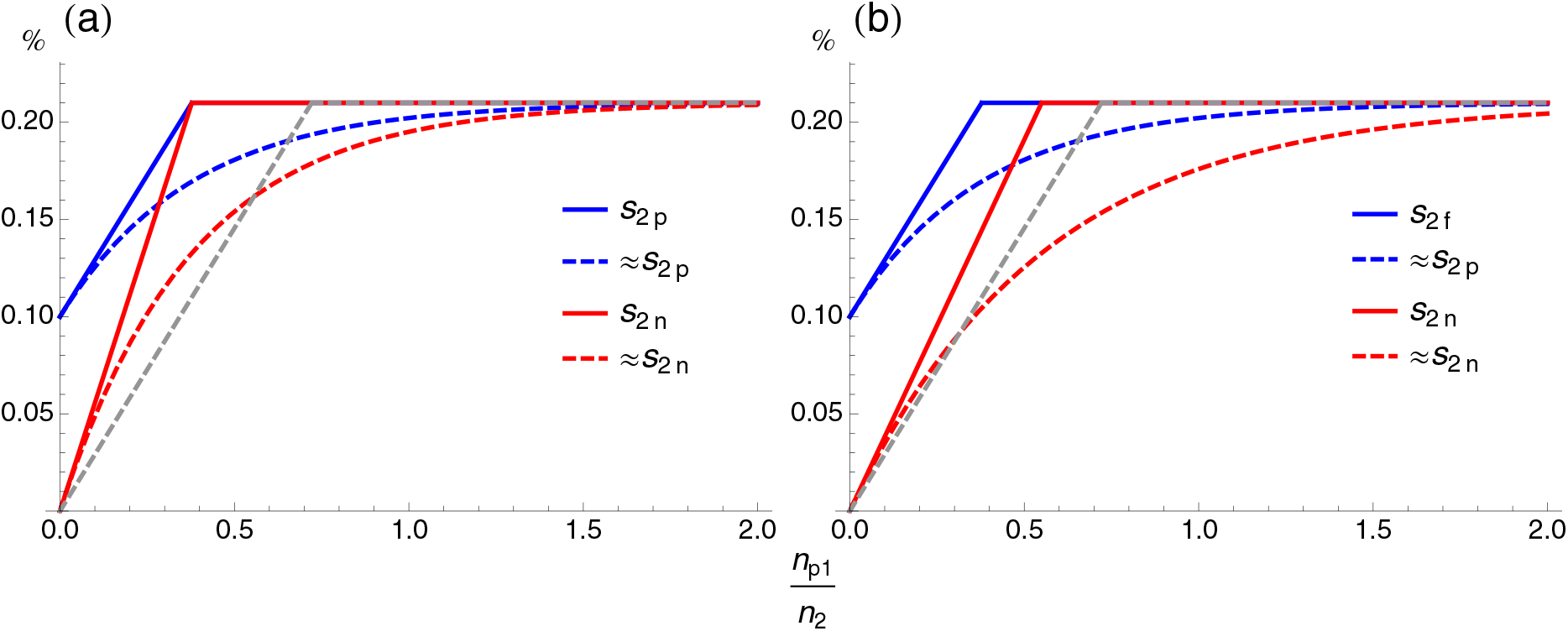
Approximations of recipients’ survival. See legend of Supporting Fig. 1. Dashed lines are the approximated survival of helped second-brood offspring that are maternally provisioned (blue) or maternally neglected (red). Such approximations were used to obtain all numerical solutions.

**Supporting Figure 3:**
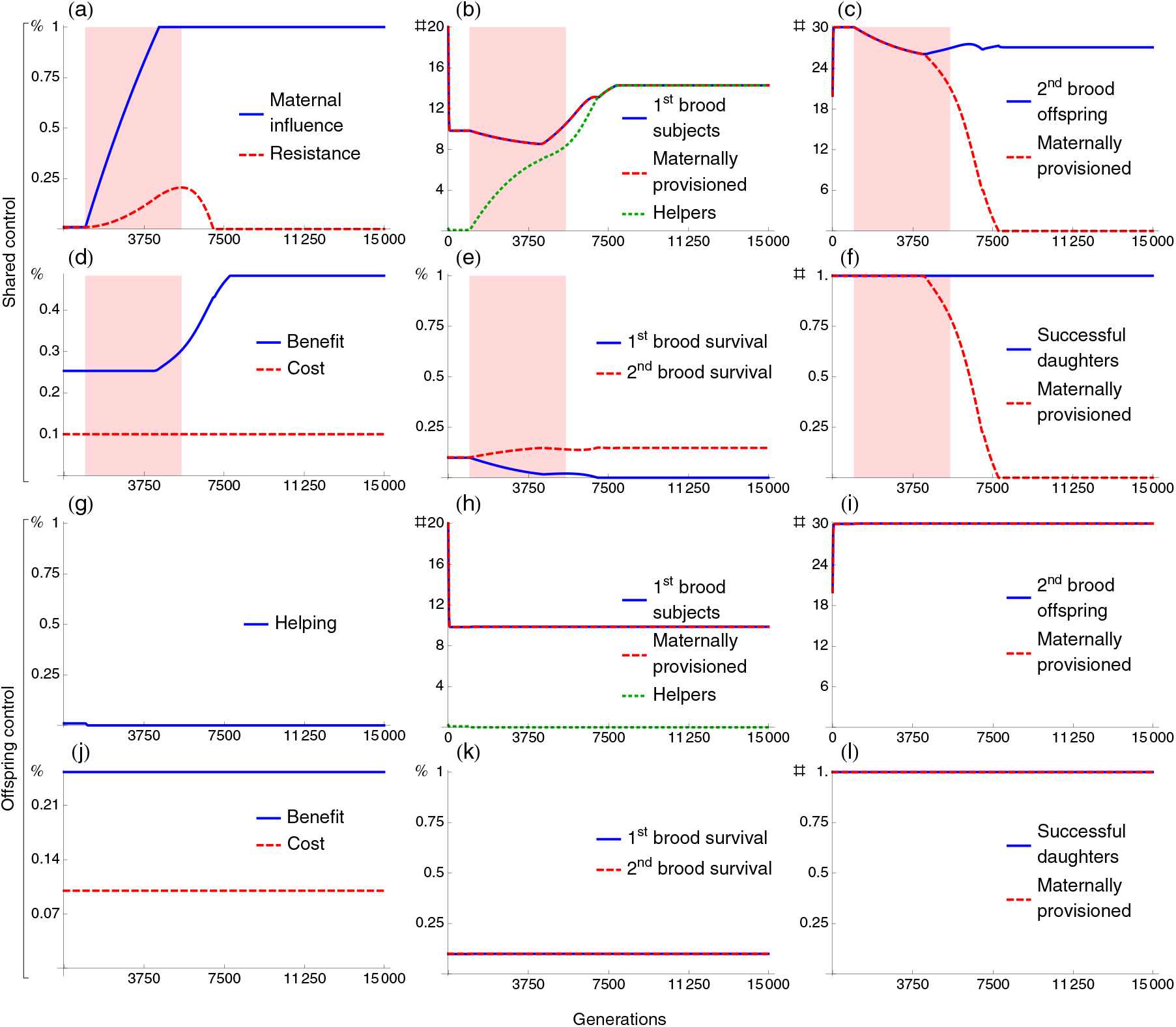
Stable eusociality via maternal manipulation with costless resistance in diploids. See legend of Fig. 2. Parameter values are in the Supporting Table 1.

**Supporting Figure 4:**
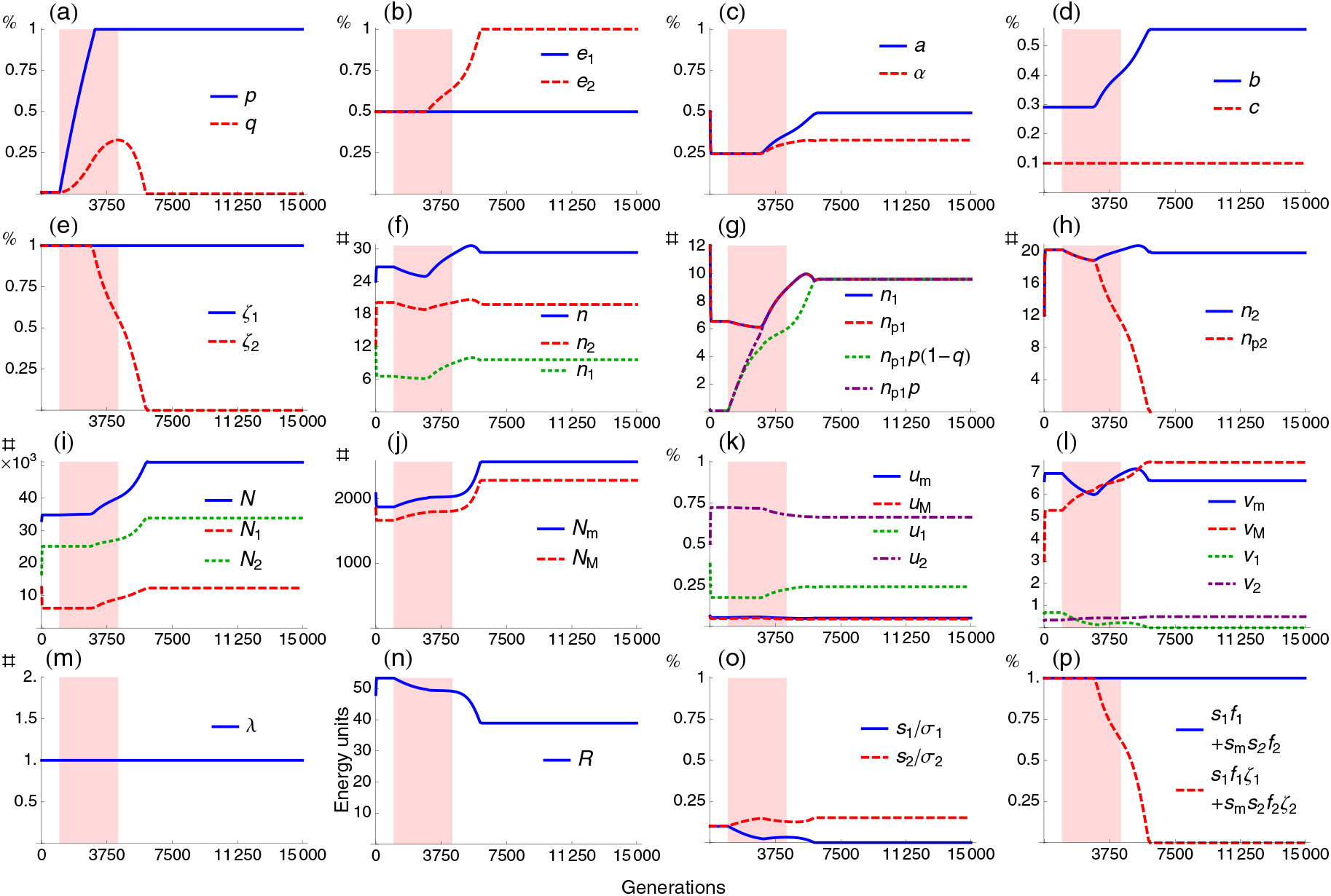
Detailed dynamics for haplodiploids under shared control. See legend of Fig. 2a-f. See Table 2 for definitions of variables. (b) The mother increases her investment in producing second-brood offspring. (h) The number of second-brood offspring remains largely constant. (i) Population size start to increase in evolutionary time when the mother increases here investment in second-brood offspring production. (m) Population size remains constant in ecological time since the ecologically asymptotic population growth rate remains 1. (n) Maternal resource decreases when the average offspring survival increases. (l) Reproductive values evolve and old mothers and second-brood offspring become more valuable. (g) *n*_p1_*p*(1 *− q*) is the number of helpers. (o) *s*_*i*_ /*σ*_*i*_ is the probability that a brood-*i* offspring becomes a parent. (p) *s*_1_ *f*_1_ *+s*_m_*s*_2_ *f*_2_ is the number of daughters that become mothers weighted by maternal genetic contribution. *s*_1_ *f*_1_*ζ*_1_ *+ s*_m_*s*_2_ *f*_2_*ζ*_2_ is the number of them that are maternally provisioned.

**Supporting Figure 5:**
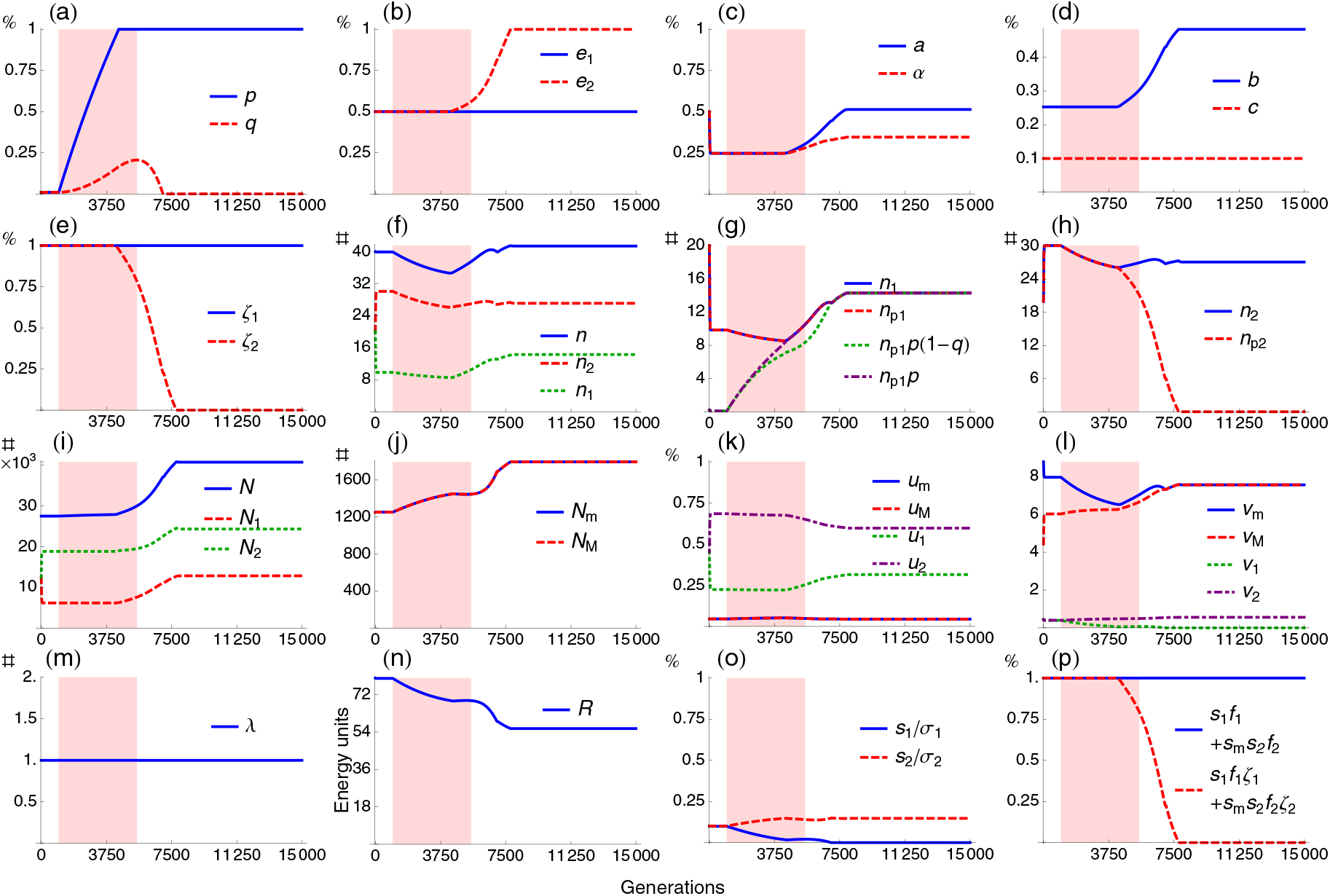
Detailed dynamics for diploids under shared control. See legend of Supporting Figs. 3a-f and 4.

**Supporting Figure 6:**
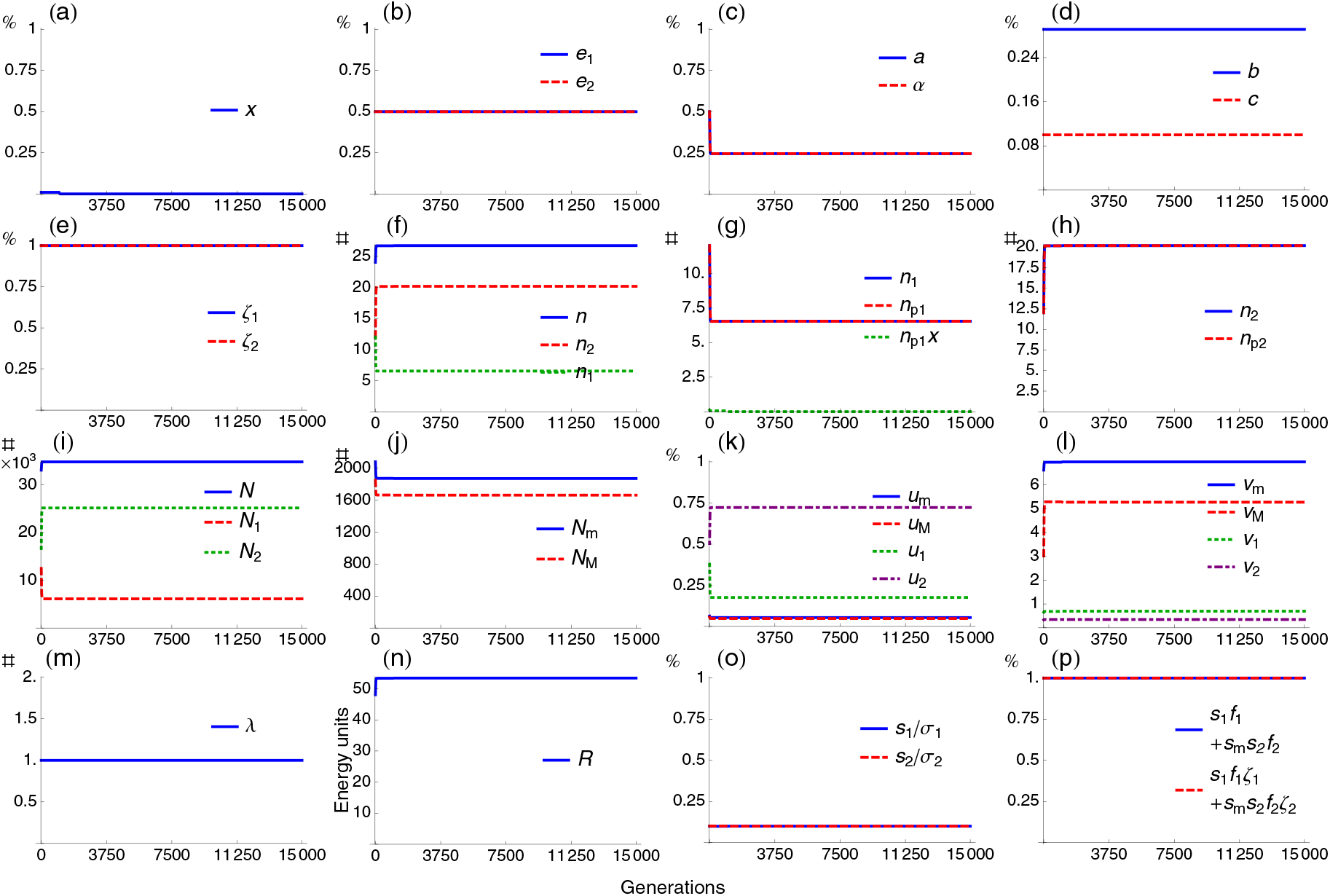
Detailed dynamics for haplodiploids under offspring control. See legend of Fig. 2g-l and Supporting Fig. 4. (a) *x* is the population-average probability that a first-brood subject stays in the natal nest in the absence of maternal influence.

**Supporting Figure 7:**
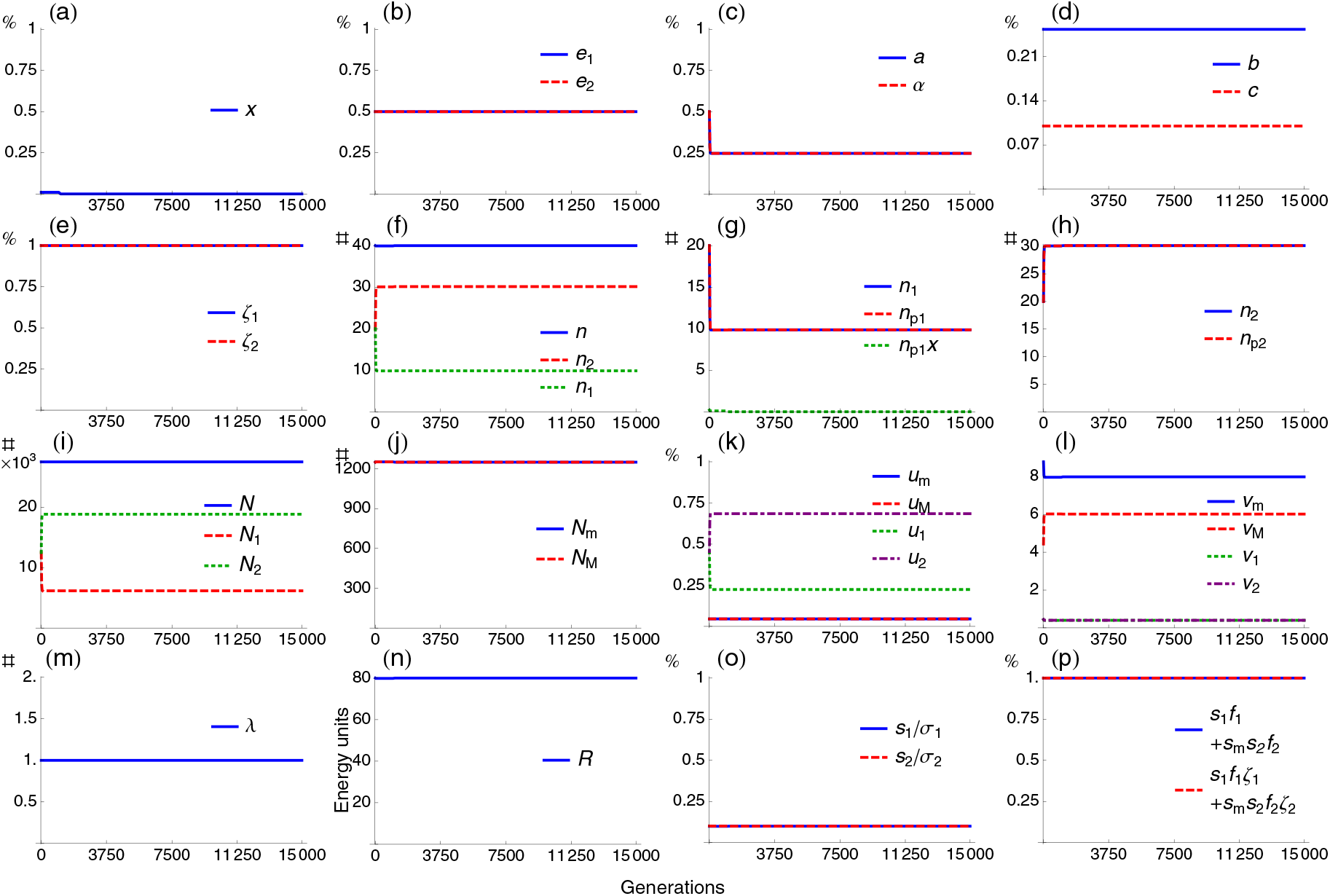
Detailed dynamics for diploids under offspring control. See legend of Supporting Figs. 3g-l and 6.

**Supporting Figure 9-14:**
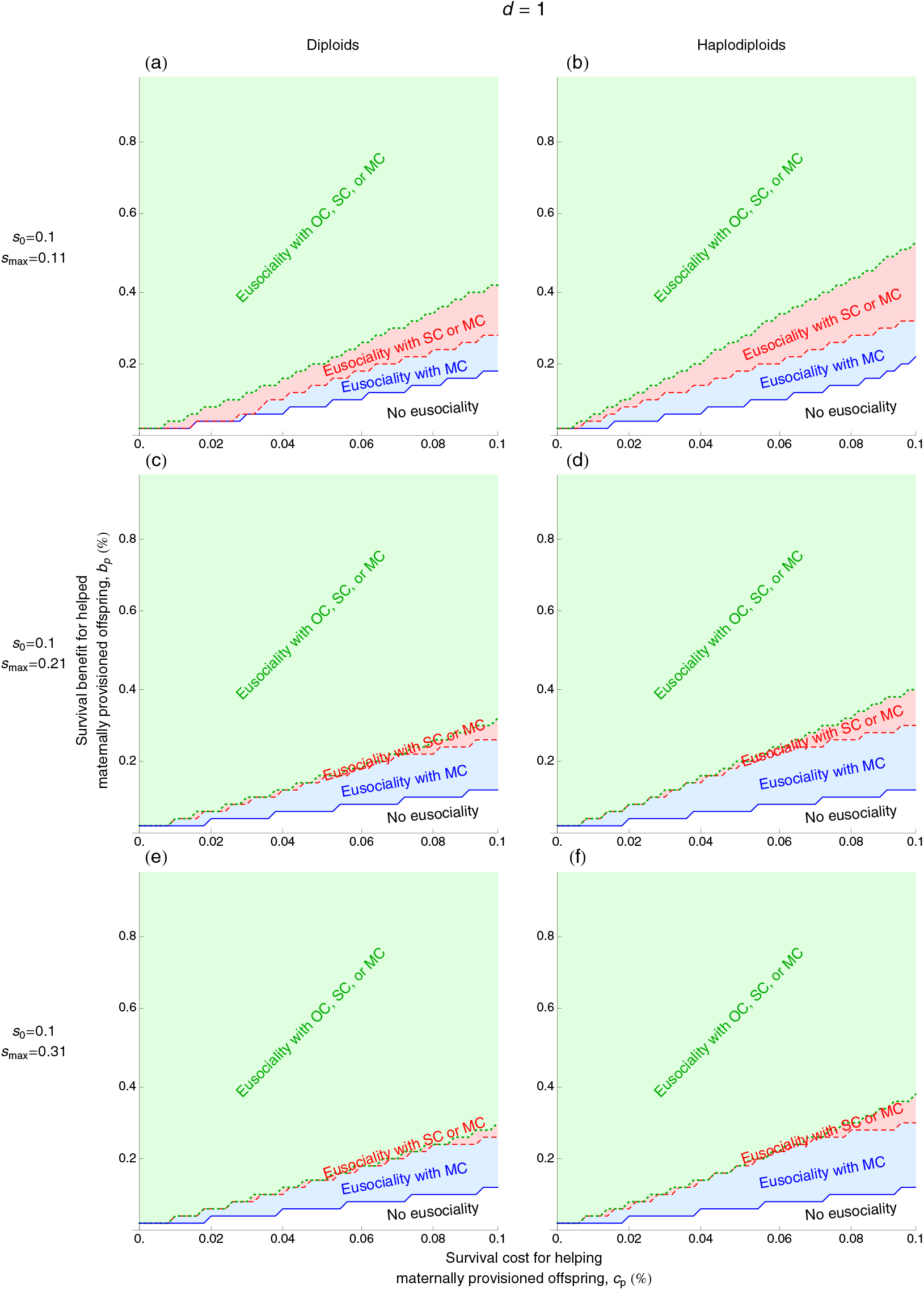
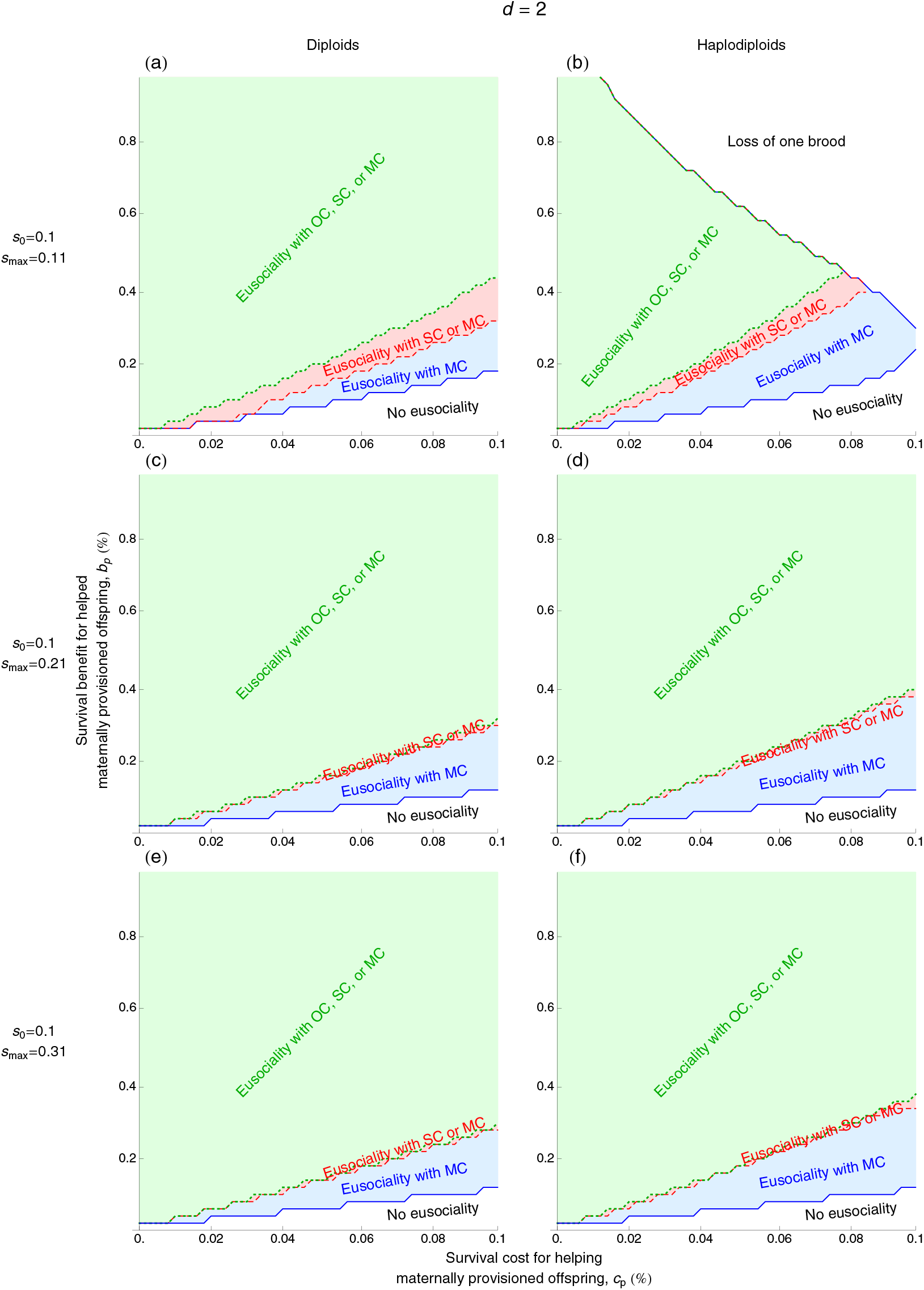
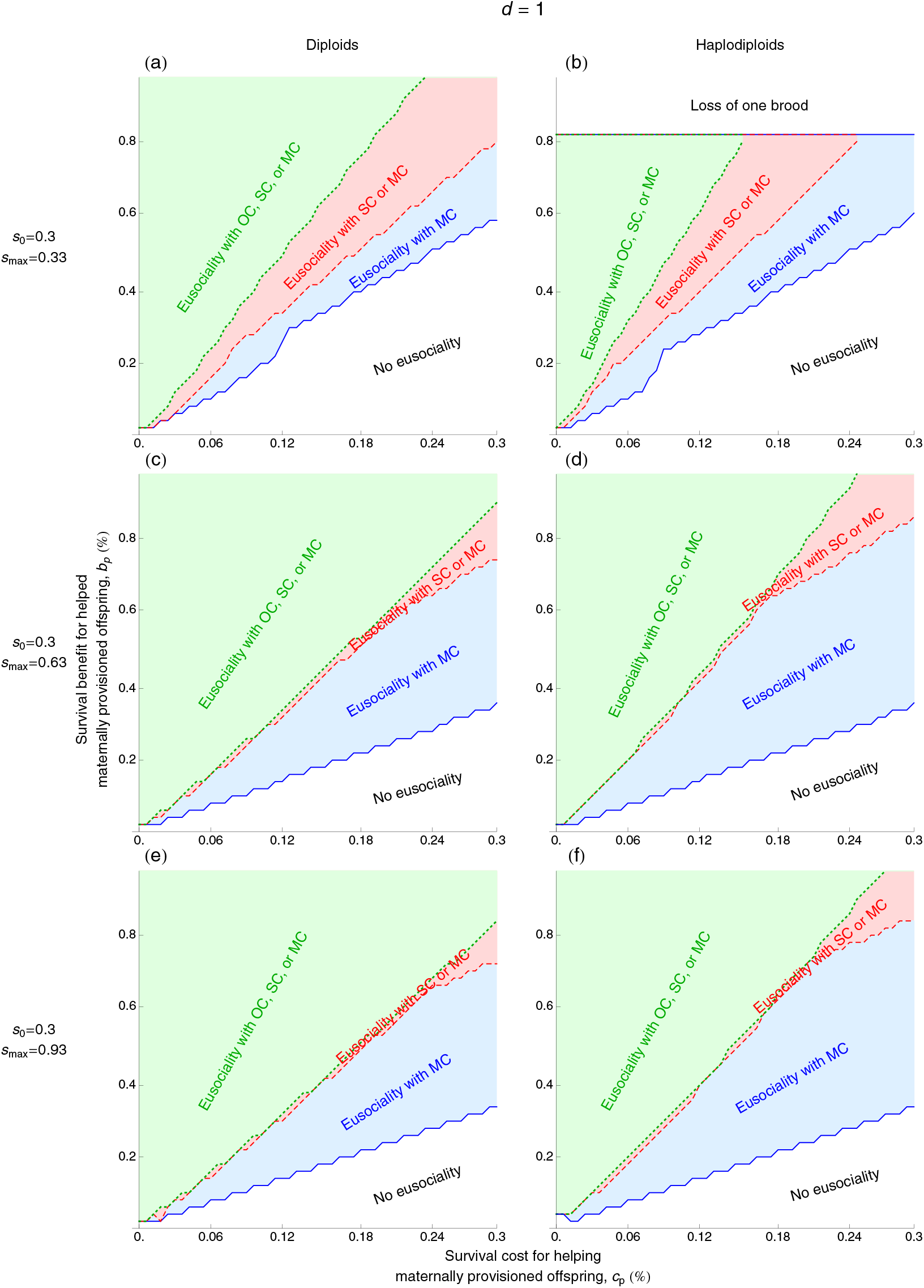
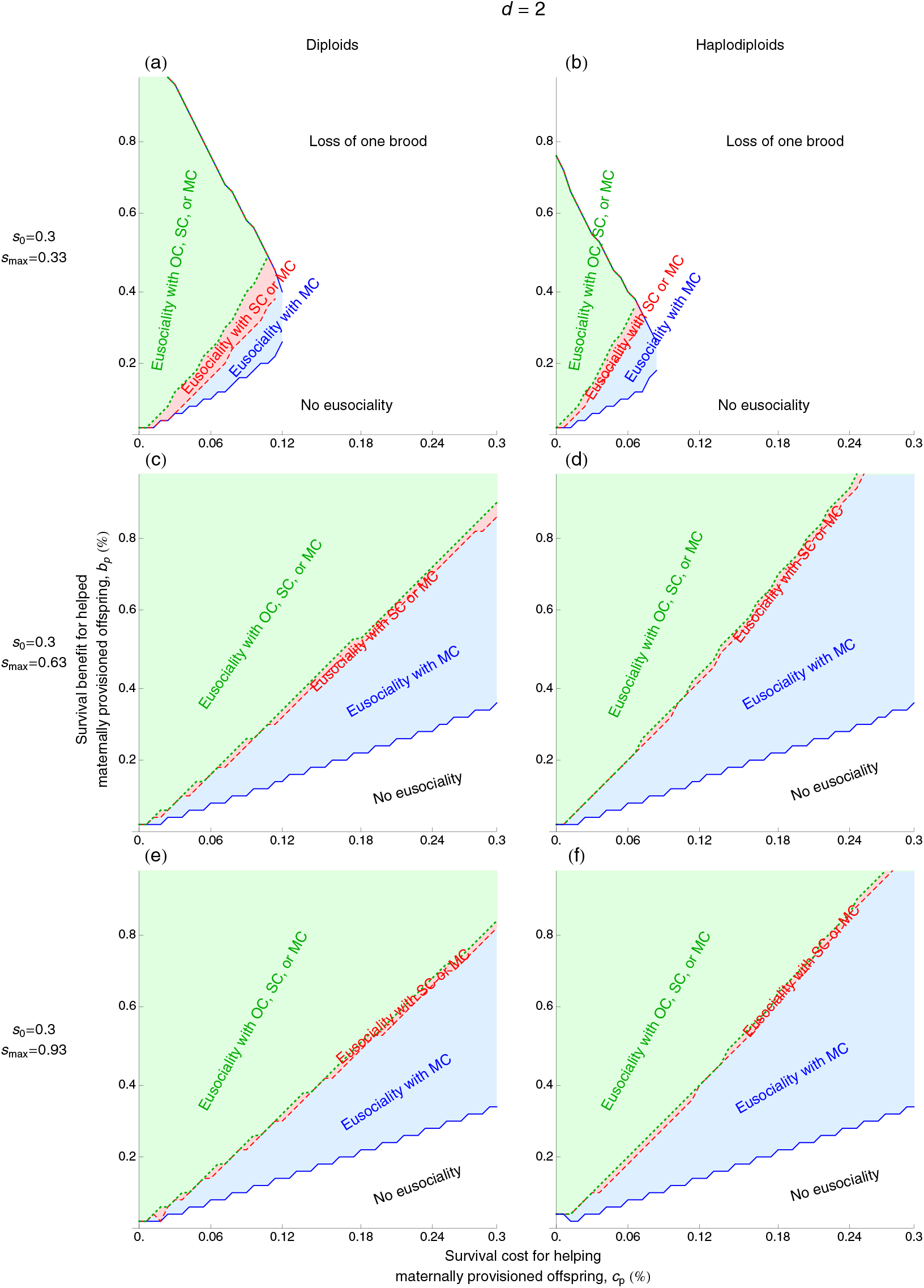
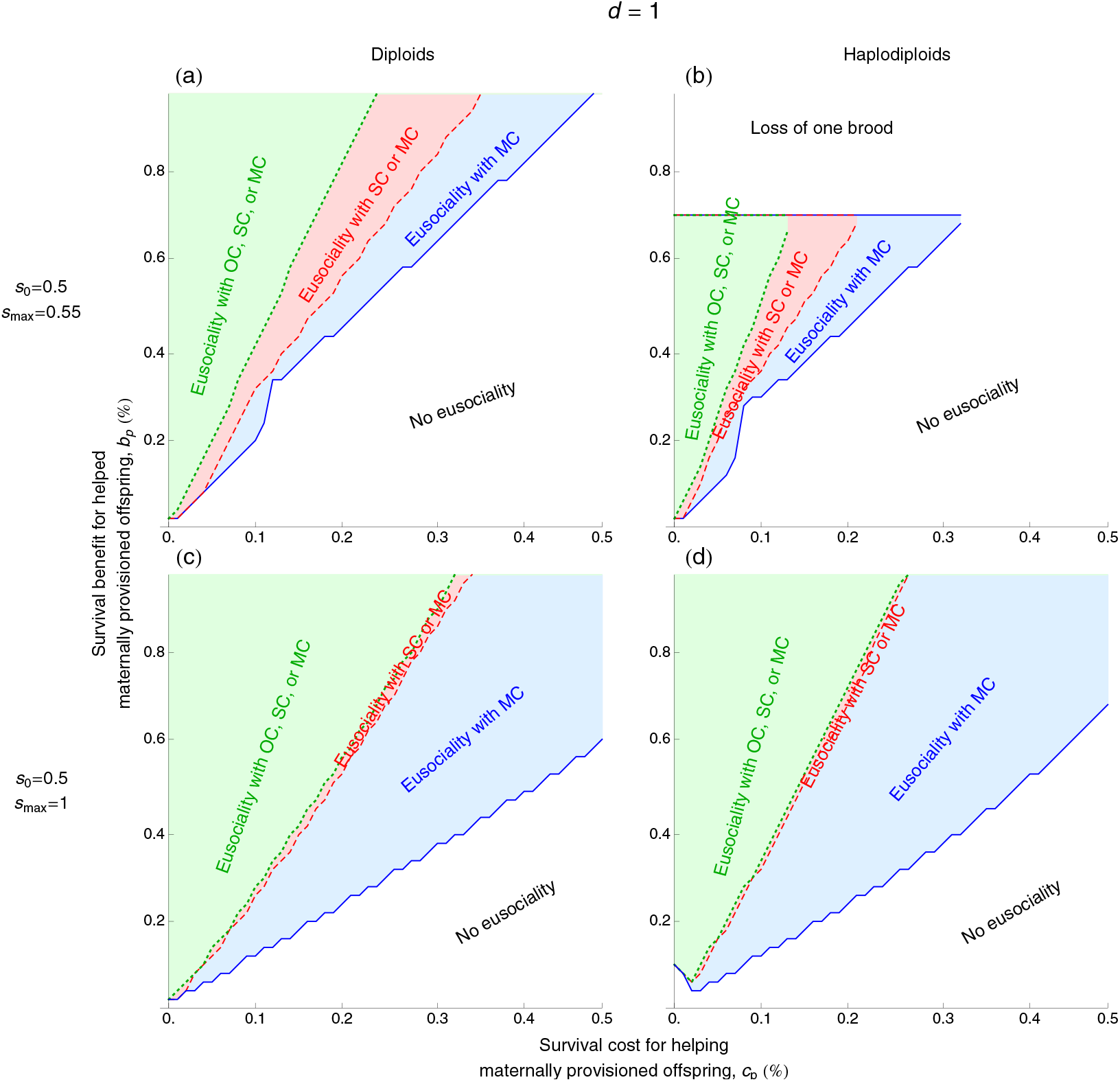
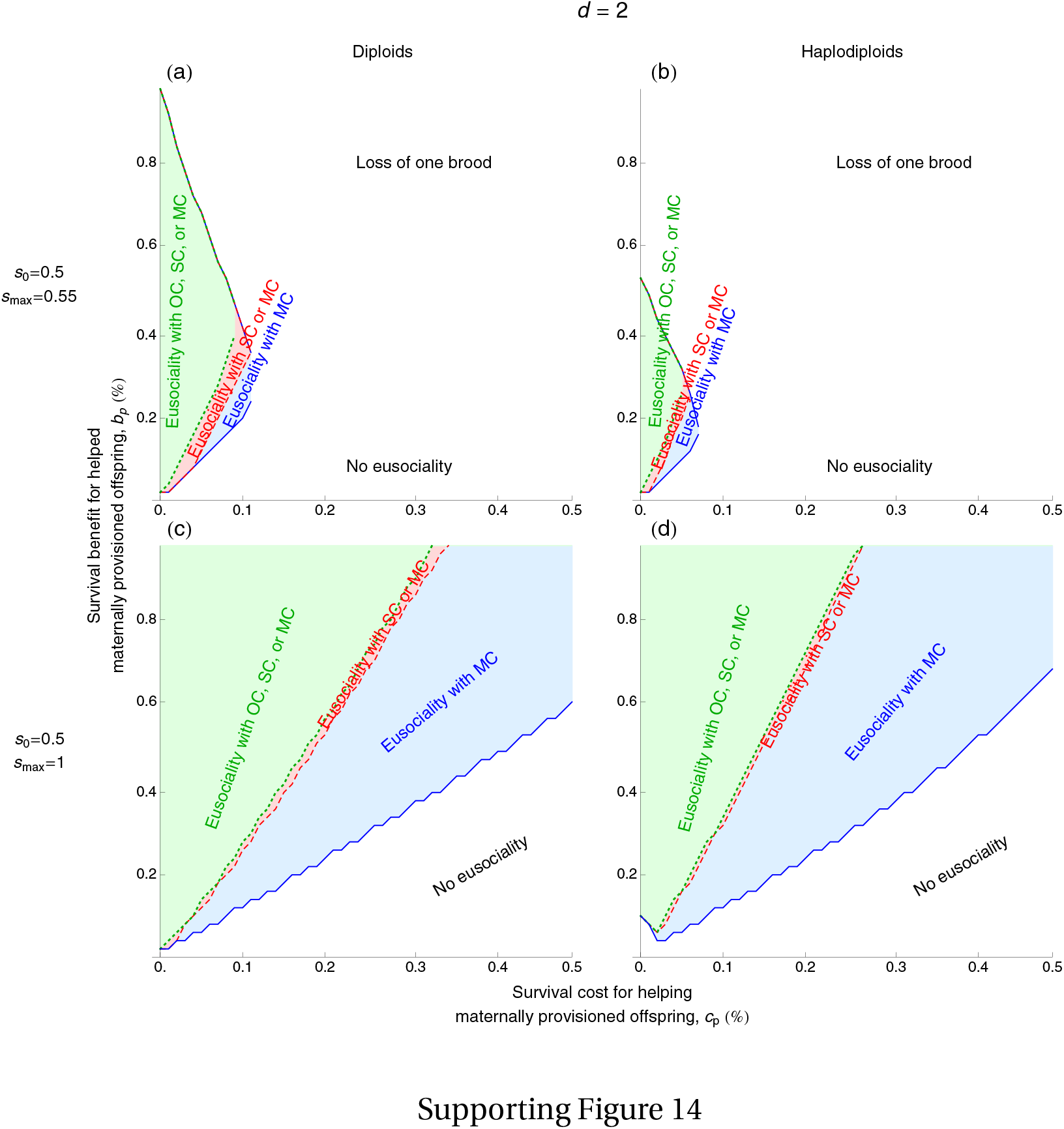
Parameter space exploration. See legend of Fig. 3 in the main text. Baseline survival is small (*s*_0_ *=* 0.1) in Supporting Figs. 9 and 10; intermediate (*s*_0_ *=* 0.3) in Supporting Figs. 11 and 12, and large (*s*_0_ *=* 0.5) in Supporting Figs. 13 and 14. The advantage of maternally neglected offspring in help use efficiency is strong (*d =* 1) for Supporting Figs. 9, 11, and 13; and weak (*d =* 2) for Supporting Figs. 10, 12, and 14. For certain regions, one of the broods is absent in the end (*n*_*i*_ *<* 1) as the mother devotes most of her resources toward one of them (Supporting Figs. 10b, 11b, 12a,b, 13b, and 14a,b; bordering lines with no eusociality are not shown). The remaining parameter values are in Supporting Table 1.

